# Gut bacterial nutrient preferences quantified in vivo

**DOI:** 10.1101/2022.01.25.477736

**Authors:** Xianfeng Zeng, Xi Xing, Meera Gupta, Felix C Keber, Jaime G Lopez, Asael Roichman, Lin Wang, Michael D Neinast, Mohamed S Donia, Martin Wühr, Cholsoon Jang, Joshua D Rabinowitz

## Abstract

Great progress has been made in understanding gut microbiome’s products and their effects on health and disease. Less attention, however, has been given to the inputs that gut bacteria consume. Here we quantitatively examine inputs and outputs of the mouse gut microbiome, using isotope tracing. The main input to microbial carbohydrate fermentation is dietary fiber, and to branched-chain fatty acids and aromatic metabolites is dietary protein. In addition, circulating host lactate, 3-hydroxybutyrate and urea (but not glucose or amino acids) feed the gut microbiome. To determine nutrient preferences across bacteria, we traced into genus-specific bacterial protein sequences. We find systematic differences in nutrient use: Most genera in the phylum Firmicutes prefer dietary protein, *Bacteroides* dietary fiber, and *Akkermansia* circulating host lactate. Such preferences correlate with microbiome composition changes in response to dietary modifications. Thus, diet shapes the microbiome by promoting the growth of bacteria that preferentially use the ingested nutrients.

## INTRODUCTION

The gut microbiome possesses an enormous diversity of enzymes, exceeding the number in mammals’ genomes by more than 100-fold (Qin et al., 2010). This enzymatic capacity enables the processing of incoming dietary nutrients into a broad spectrum of microbial metabolites. Some of these reach the host circulation at substantial concentrations (Lai et al., 2021; Quinn et al., 2020). Microbial metabolites can play important roles in host pathophysiology. For example, short-chain fatty acids (SCFAs; acetate, propionate, butyrate) (Dalile et al., 2019; Koh et al., 2016), trimethylamine N-oxide (Tang et al., 2013), secondary bile acids (Arab et al., 2017; Funabashi et al., 2020), indole-3-propionate (Wikoff et al., 2009), and imidazole propionate (Koh et al., 2018) affect immune maturation (Campbell et al., 2020; Hang et al., 2019), insulin sensitivity (Koh et al., 2018), cancer growth (Garrett, 2015; Yoshimoto et al., 2013), and cardiovascular disease (Nemet et al., 2020; Wang et al., 2011).

Both to replicate themselves and to release metabolic products, gut bacteria require nutrient inputs. These come in forms including ingested food, host-synthesized gut mucus (Desai et al., 2016; Sicard et al., 2017), and host circulating metabolites (Scheiman et al., 2019). The availability of dietary nutrients to gut microbiota depends on the extent of host absorption: nutrients that are absorbed in the small intestine, like starch, are not available to the colonic microbiome. In contrast, nutrients that are poorly digested in the upper gastrointestinal tract, like fiber, can be key microbiome feedstocks (Lund et al., 2021; Wong and Jenkins, 2007).

Isotope tracing enables the measurement of the inputs to metabolites and biomass in a quantitative manner. Such studies, employing radioactive tracers, defined the basics of mammalian metabolism (Wolfe, 1984). Recent work has increasingly relied on stable isotope tracers coupled to mass spectrometry detection, which enables the measurement of labeling in specific downstream products (Fernández-García et al., 2020; McCabe and Previs, 2004). This approach has revealed fundamental features of host metabolism, such as circulating lactate being a major TCA fuel (Faubert et al., 2017; Hui et al., 2017). In addition, it has provided important insights into host-microbiome metabolic interplay. For example, it revealed that dietary fructose is processed by the microbiome into acetate, which fuels hepatic lipogenesis (Jang et al., 2018; Zhao et al., 2020).

In principle, stable isotope tracing coupled to mass spectrometry can also be applied to determine the metabolic inputs to specific microbes, based on measuring labeling in bacteria-specific peptide sequences (Berry et al., 2015; Holmes et al., 2017; Oberbach et al., 2017; Reese et al., 2018; Zhang et al., 2016a, 2016b). By infusing nitrogen-labeled threonine to label host mucus, investigators were able to compare the contribution of dietary versus mucus protein to the gut microbiome and observed a shift towards more mucus contribution in a low-protein diet condition (Holmes et al., 2017).

Here, we perform the first large-scale, quantitative assessment of the metabolic inputs to the gut microbiome and its products. We examine the contributions from dietary starch, fiber, and protein, host mucus, and most major circulating host nutrients, finding that lactate, 3-hydroxybutyrate, and urea stand out for passing from the host to the gut microbiome. Moreover, based on the measurement of bacteria-specific peptide sequences, we assess the nutrient preferences of different bacterial genera and show that these preferences align with microbiome composition changes in response to altered diet.

## RESULTS

### Microbiome consumes less digestible dietary components

A major mechanism by which the microbiome may impact host physiology is via secreted metabolic products. As the intestine and colon drain into the portal circulation, metabolites produced by the gut microbiome should be enriched in the portal relative to systemic blood. We measured, in the portal, systemic circulation and the cecal contents, the absolute concentrations of more than 50 metabolites characterized in the literature as microbiome-derived (Campbell et al., 2020; De Vadder et al., 2014; Han et al., 2021; Hang et al., 2019; Koh et al., 2018; Mager et al., 2020; Ridlon et al., 2014; Wikoff et al., 2009) (**Figure 1A, S1A, Table 1, S1-2**). Most microbiome metabolites were elevated in the portal circulation relative to systemic blood, and all but two (inosine and N-acetyl-tryptophan, which are apparently mainly host derived) were depleted by antibiotics treatment.

**Table 1.**
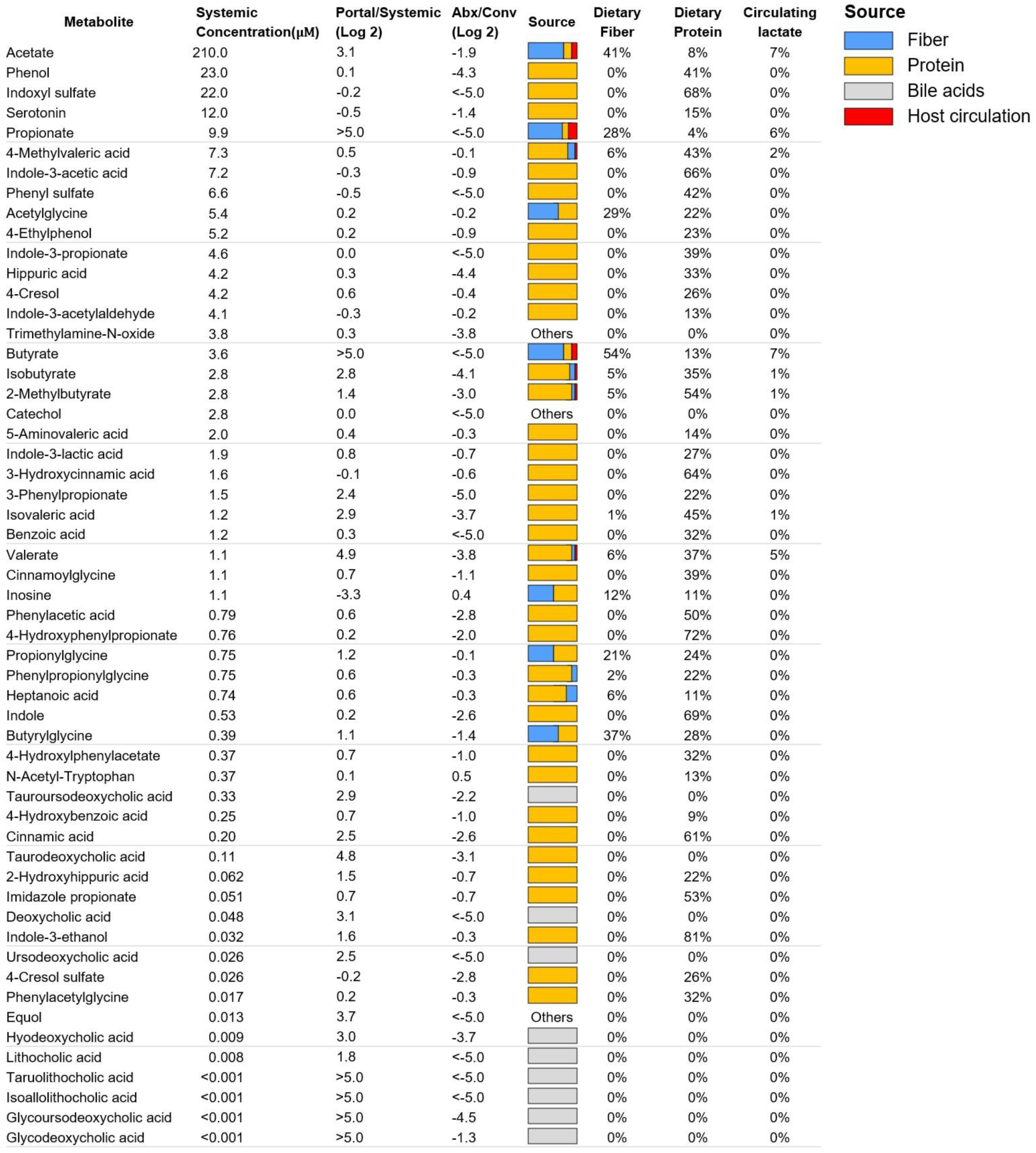
Absolute concentrations and sources of microbiota-associated metabolites. Data are from ad lib fed state (ZT0); for ad lib fasted state (ZT12), see Supplementary Table S1. Absolute concentration is mean, N = 5 mice. Portal/systemic = fold-change in concentration between the portal vein and tail vein (median, N = 5 mice). Abx/Conv refers to fold-change in portal blood concentration between mice treated with antibiotics cocktail versus not (median, N = 5 mice/group). Source bar indicates the relative contribution to the indicated metabolite from dietary inulin, algal protein and circulating lactate (based on isotope tracing). Percentages indicate quantitative relative contributions from those nutrients  (median, N = 4). Numbers typically add up to less than 100%, as other sources (e.g. mucins) contribute.

**Figure 1.**
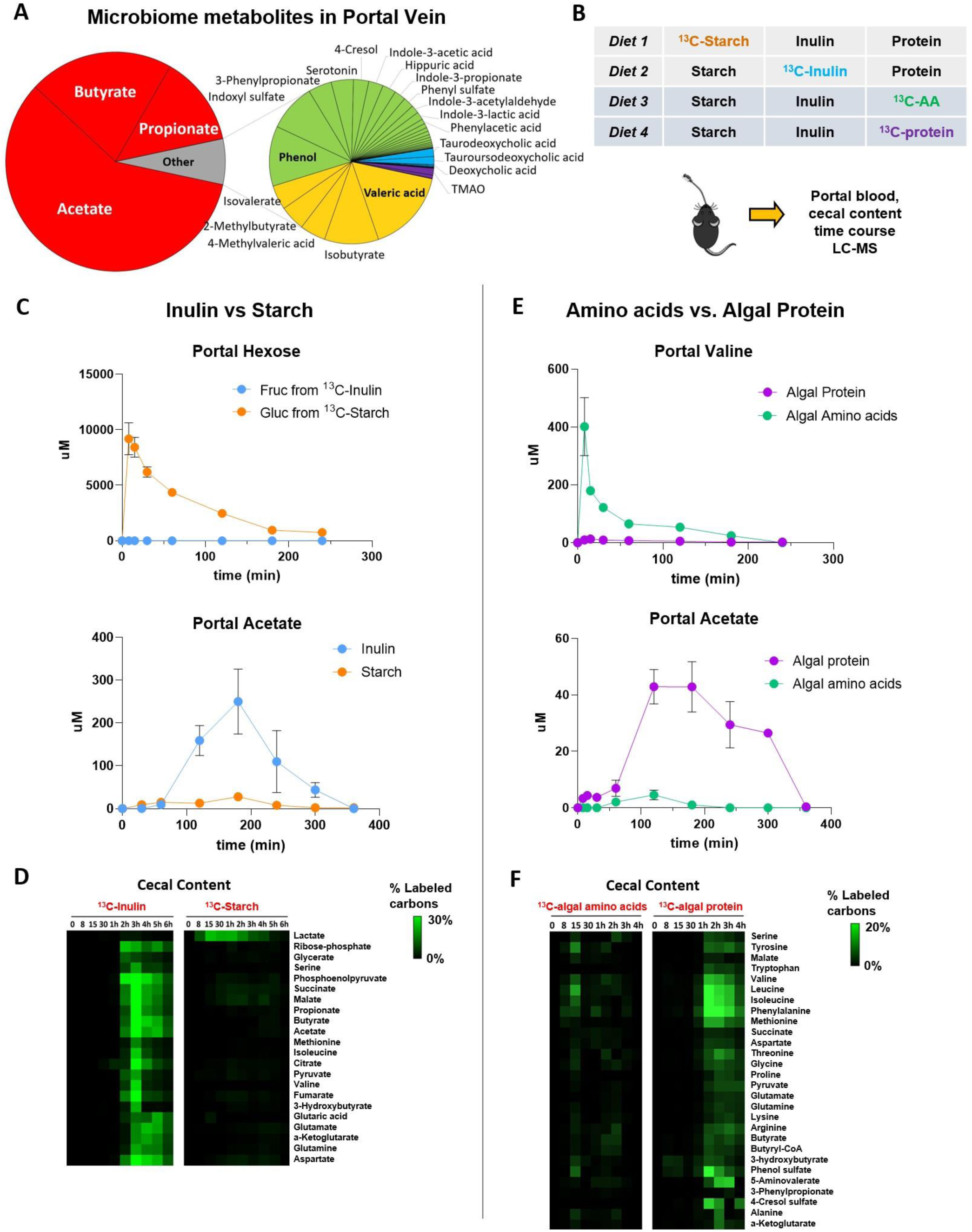
Microbiome consumes dietary fiber and protein. (A) Composition of the measured portal microbial metabolome. The pie charts show the relative molar abundance of different gut microbiota-associated metabolites in mice (N = 6 mice). (B) Experimental scheme. Mice received an oral gavage of 4:2:1 starch: protein (or free amino acids): inulin by weight. In each dietary condition, one component was ^13^C-labeled. After gavage of the labeled diet, tissue and serum metabolite labeling were measured over time by LC-MS. (C) Dietary starch feeds the host, while dietary inulin feeds the microbiome. The data shows concentrations of labeled carbons in hexose and acetate in portal circulation (mean ± s.e., N = 3 mice). (D) Inulin is a major microbiome feedstock. Heatmap shows the percentage of labeled carbon atoms in the indicated metabolites in cecal content. Each data point is median of N = 3 mice. (E) Dietary amino acids feed the host while dietary algal protein feeds the microbiome. The data shows concentrations of labeled carbons in valine and acetate in portal circulation (mean ± s.e., N = 3 mice). (F) Protein but not free amino acids are a major microbiome feedstock. Heatmap shows the percentage of labeled carbon atoms in the indicated metabolites in cecal content. Each data point is median of N = 3 mice.

The dominant excreted products on a molar basis (0.4 – 2 mM in the portal blood) are SCFAs. Other relatively abundant microbiome products (10 - 30 uM) are aromatic amino acid fermentation products (phenol, indoxyl sulfate, and 3-phenylpropionate) and branched-chain fatty acids (valerate, isovalerate, 4-methylvalerate, isobutyrate, 2-methylbutyrate). While primary bile acids were also present in the portal circulation at up to ∼ 10 uM concentration, these are produced by the host and accordingly were not included in Table 1. Secondary bile acids, which are produced from primary bile acids by the microbiome, were lower in absolute concentration, the most abundant being tauroursodeoxycholic acid (3 uM in portal vein).

To probe the dietary inputs to gut microbial products, we began by feeding mice, by oral gavage, starch (readily digestible glucose polymer) and inulin (slowly digestible fructose polymer, i.e., soluble fiber) (**Figure 1B**). Following ^13^C-starch gavage, labeled glucose, lactate, and alanine quickly appeared in the portal circulation (**Figure 1C, S1B**). In contrast, after ^13^C-inulin gavage, substantial labeled fructose, glucose, lactate, and alanine were not observed, and instead labeled portal metabolites slowly appeared in the form of SCFAs (**Figure 1C, S1C**). Quantitative analysis based on the portal-systemic differences in serum metabolite labeling (Jang et al., 2018) revealed that most starch carbons (∼75%) become circulating glucose, lactate, and alanine, with no discernible labeling of SCFAs. In contrast, ∼ 40% of inulin carbons become SCFAs (**Figure S1C**), with the remainder being undigested and excreted in the feces (**Figure S1D**). Measurement of cecal content revealed that dietary inulin, but not starch, extensively labeled glycolytic and TCA intermediates and amino acids in the cecal content (**Figure 1D**).

We next carried out similar experiments, comparing the gavage of a free amino acid mixture to algal protein, both uniformly ^13^C-labeled (**Figure 1B**). The free amino acid feeding resulted in the rapid appearance of labeled amino acids in portal circulation, while the algal protein did not (**Figure 1E**). Instead, the algal protein, but not free amino acids, substantially labeled amino acids within the cecal contents (**Figure 1F**). Moreover, the algal protein copiously labeled microbiome-derived portal vein metabolites: SCFAs, branched-chain fatty acids, and aromatics (indole, indole-3-propionate, 3-phenylpropionate) (**Figure S1E-F**). Thus, poorly digestible carbohydrates and protein feed the microbiome directly, and the host indirectly via microbiome-derived products.

### Few circulating metabolites reach the microbiome

Next, we examined the possibility that nutrients in host circulation feed the gut microbiota. We infused deuterated water and eighteen major circulating nutrients (^13^C-labeled) into the systemic circulation of pre-catheterized mice (**Figure 2A**). The infusion rates were selected to achieve modest but readily measurable labeling without substantially perturbing circulating concentrations. Circulating labeling reached a steady-state by 2.5 h, at which time we collected serum and feces to quantitate the carbon contributions of each circulating nutrient to the corresponding fecal metabolites. Upon intravenous infusion of ^13^C-lactate, fecal lactate labeled rapidly (**Figure 2B**). Most infused circulating nutrients, however, did not penetrate the feces (**Figure 2C-D**). Indeed, while water fully exchanged with the feces, among abundant circulating carbon carriers, only lactate and 3-hydroxybutyrate penetrated. Glucose, amino acids, TCA intermediates and fatty acids did not. Both lactate and 3-hydroxybutyrate are substrates of monocarboxylate transporters (MCTs), which are highly expressed in the colonic epithelium (Halestrap and Price, 1999, p. 1). Thus, in contrast to most host circulating metabolites, which do not reach the colonic microbiome, monocarboxylic transporters render circulating lactate and 3-hydroxybutyrate accessible to gut microbes.

**Figure 2.**
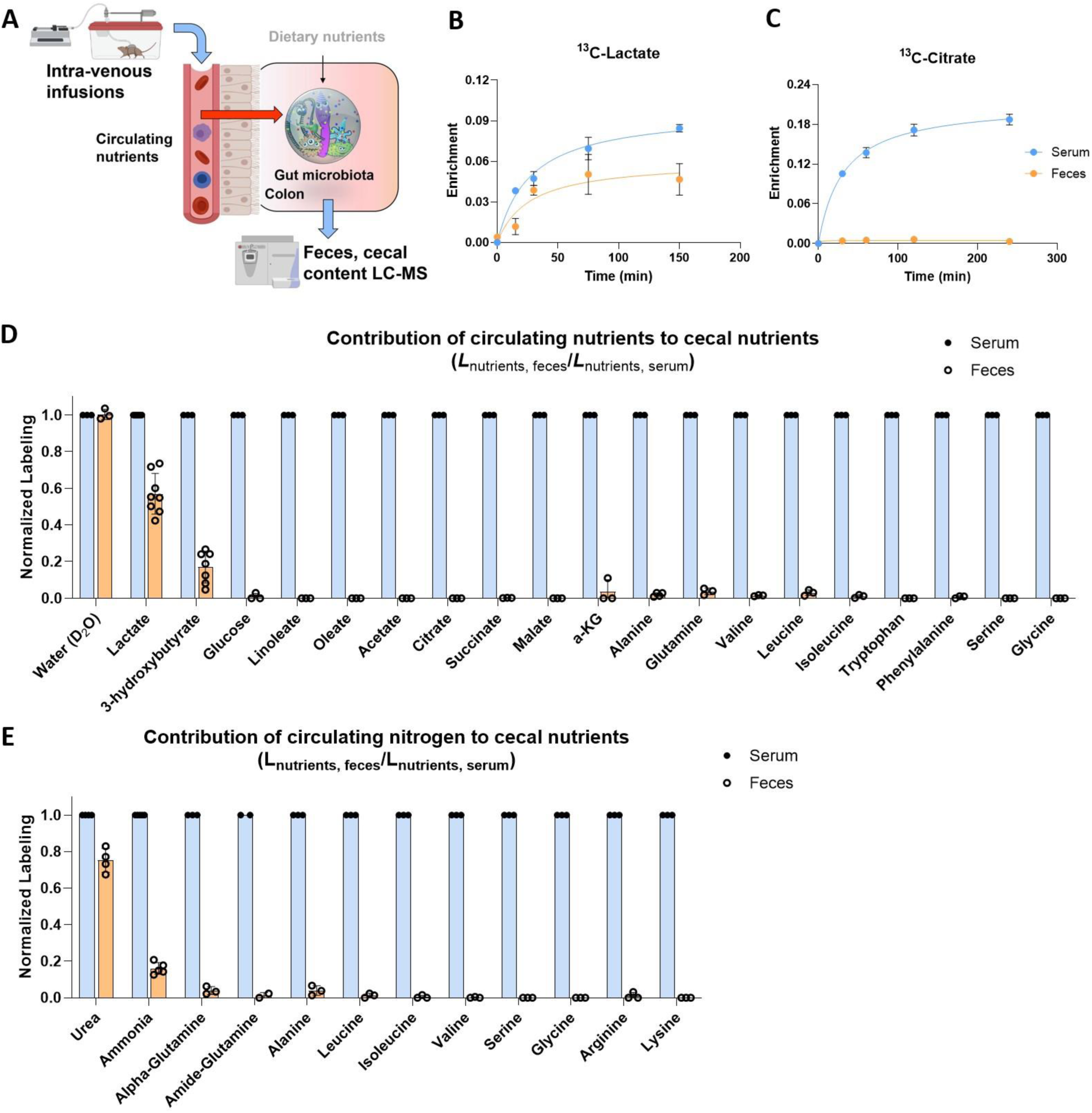
Circulating lactate and 3-hydroxybutyrate feed the gut microbiome. (A) Schematic of intravenous infusion of isotope-labeled nutrients to identify circulating metabolites that feed gut microbiome. (B) Circulating lactate rapidly enters the feces. Mice were infused with ^13^C-lactate and serum and fresh feces enrichment were compared. Mean±s.e. N = 3. (C) Circulating citrate does not enter the feces. As in (B), for ^13^C-citrate. (D) Passage of circulating ^13^C-labeled nutrients into the feces. Mice were infused with labeled nutrients for 2.5 h, and labeling fraction in feces was normalized to labeling fraction in serum. Mean±s.e. N = 3 except for lactate (N = 8) and 3-hydroxybutyrate (N = 7). (E) Passage of circulating ^15^N-labeled nutrients into the feces. As in (D), for ^15^N-lableing. Mean±s.e. N = 3 except for urea (N = 4) and ammonia (N = 5).

### Circulating urea is a microbiome nitrogen source

In addition to carbon, nitrogen is a fundamental constituent of all living cells. To assess nitrogen sources of the gut microbiome, we infused twelve abundant circulating nutrients in ^15^N-labeled form. Nitrogen from circulating urea and ammonia, but not amino acids, penetrates the feces and contributes to microbiome amino acids (**Figure 2E, S2A-B**).

The liver converts ammonia into urea. Accordingly, we were curious if ammonia contributes to the microbiome directly, or only indirectly after being converted by the host into circulating urea (Bartman et al., 2021). Ammonia’s contribution is quantitatively explained by the product of its contribution to circulating urea and urea’s microbiome contribution (**Figure S2C-D**). Thus, urea is the main circulating host metabolite that provides nitrogen to the gut microbiome.

### Microbiota synthesize amino acids from fiber and urea

To determine the physiological sources of microbiome metabolites, we measured their labeling after *ad libitum* feeding of isotopically enriched food. To this end, we fed mice standard chow with a portion of the fiber, fat, or protein ^13^C-labeled. To account for circulating nutrient inputs, we also infused ^13^C-lactate or 3-hydroxybutyrate (**Figure 3A, S3A**). These studies identified a majority of the carbon feeding into most microbiome central metabolites, with glycolytic and pentose phosphate metabolites labeling almost exclusively coming from dietary fiber (inulin), while pyruvate and TCA metabolites are also labeled from dietary algal protein and circulating lactate (**Figure 3B**).

**Figure 3.**
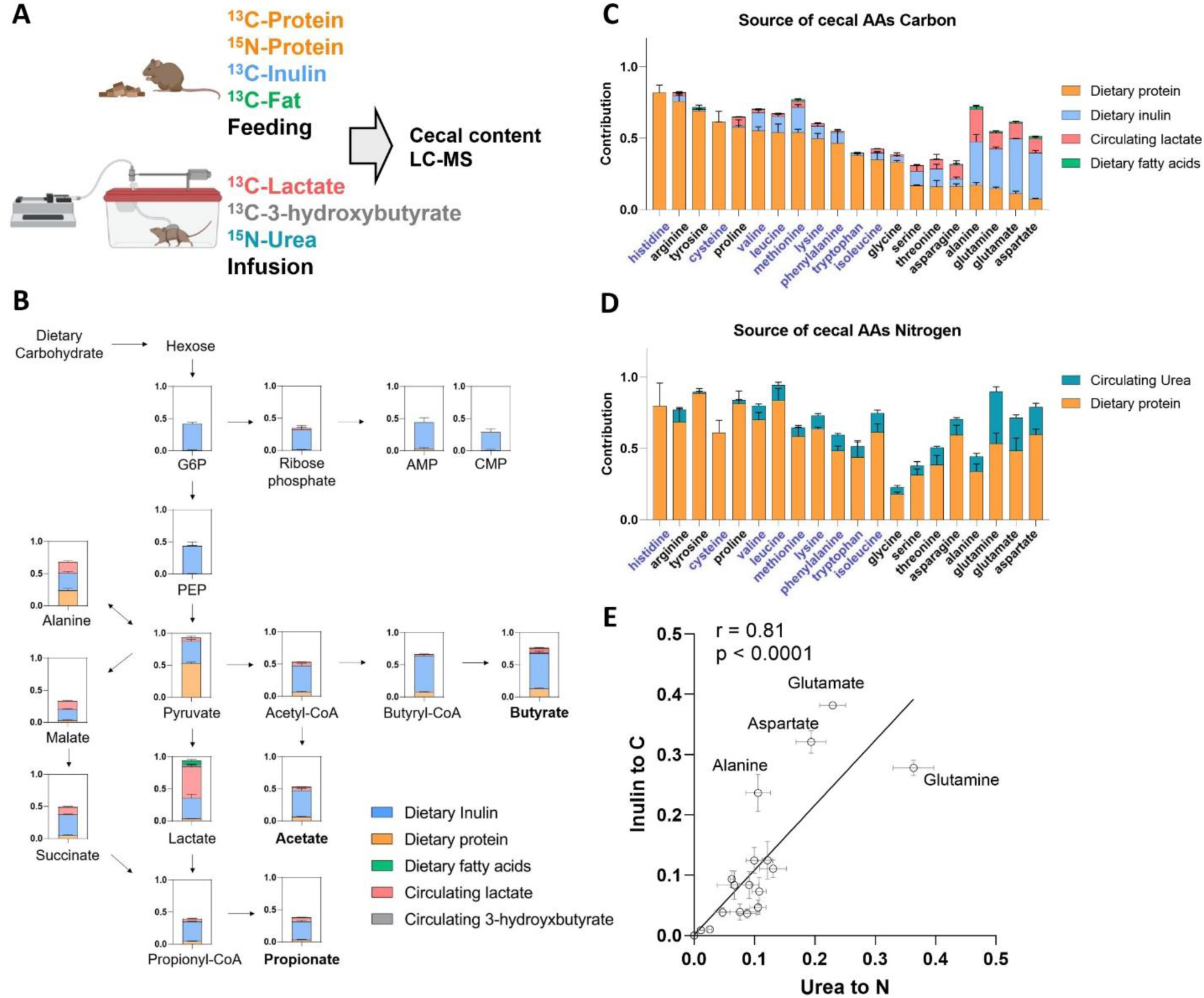
Quantitative analysis of dietary and circulating nutrient contributions to gut microbiome. (A) Experimental design. Mice were fed chow containing ^13^C-protein, ^13^C-inulin, ^13^C-fatty acids, or ^15^N-protein for 24 h. Alternatively, mice were intravenously infused with ^13^C-lactate, ^13^C-3-hydroxybutyrate or ^15^N-urea for 24 h. The labeling of cecal content metabolites was analyzed by LC-MS. (B) Contribution of dietary and circulating nutrients to carbohydrate fermentation pathways in gut microbiome. Mean ± s.e. N = 4. (C) Contribution of dietary and circulating nutrients to cecal amino acid carbon. The names of essential amino acids (EAA) are written in blue and non-essential amino acids (NEAA) in black. Mean ± s.e. N = 4. (D) Contribution of dietary and circulating nutrients to cecal amino acid nitrogen. As in (C), for nitrogen. (E) Positive correlation, across amino acids in the cecal contents, of carbon contribution from dietary inulin and nitrogen contribution from circulating urea. Mean ± s.e. N = 4.

We next examined inputs to microbiome amino acids, tracing also with ^15^N-labeled dietary protein and infused urea. Unlike mammals, most gut bacteria have the biosynthetic capacity to make all 20 proteogenic amino acids. Nevertheless, we observed that “essential amino acids,” which cannot be made by mammals and require the expression of extensive biosynthetic pathways in bacteria, are derived mainly from dietary proteins. In contrast, “non-essential amino acids” are primarily synthesized within the gut microbiome, using dietary inulin and circulating lactate as carbon sources (**Figure 3C**). Dietary protein was the main nitrogen source for both essential and non-essential amino acids, with host urea also contributing substantially to the non-essential amino acids (**Figure 3D**). Importantly, the amino acids synthesized by the microbiome, stay in the microbiome: We do not observe discernible labeling of these amino acids in the host (**Figure S3B**). Consistent with the gut microbiome synthesizing amino acids from fiber carbon and urea nitrogen, across amino acids, urea’s nitrogen contribution correlated with inulin’s carbon contribution (**Figure 3E**). Thus, the microbiome obtains amino acids from a blend of dietary protein catabolism and *de novo* synthesis fed by dietary fiber and urea.

### Diverse microbiome products come from dietary protein

We next examined, using isotope tracing, the carbon inputs to the microbiome products, especially the ones excreted into the portal circulation (**Table 1**). SCFAs, the most abundant microbial metabolites, come mainly from dietary fiber with minor contributions from dietary protein and host circulating lactate. Many less abundant ones, however, are almost exclusively derived from dietary protein (**Table 1**).

In addition to classical microbiome products, we also observed metabolites that are made in a collaborative manner, with the host carrying out the final synthesis using microbiome-derived inputs. For example, a wide range of microbiome-derived carboxylic acids are conjugated to glycine in the liver and the kidneys to make different acyl-glycines (**Figure S4A-D**) (Wikoff et al., 2009).

We also examined the host clearance mechanisms of microbiome metabolites, based on arterial-venous gradients across the liver and kidney and levels in the urine. SCFAs and branched-chain fatty acids were avidly consumed by the liver, consistent with their much greater abundance in the portal than systemic circulation. Most microbiome-derived metabolites were excreted by the kidney into the urine, with the notable exception of SCFAs, which are actively reabsorbed (**Table S1**). Thus, we establish dietary protein as a major precursor to many microbiome metabolites and identify host-microbiome interplay in the metabolism of SCFAs, including their renal reabsorption and use by liver and kidney for the synthesis of acyl-glycines.

### Gut bacterial growth is synchronized with host feeding

Thus far, we have reported inputs and outputs of the gut microbiome as a whole. We now shift to examining the growth and metabolism of specific bacterial genera. To this end, we deployed proteomics to measure gut microbial peptides and their labeling, focusing on peptide sequences specific to a single bacterial genus (**Figure 4A**).

**Figure 4.**
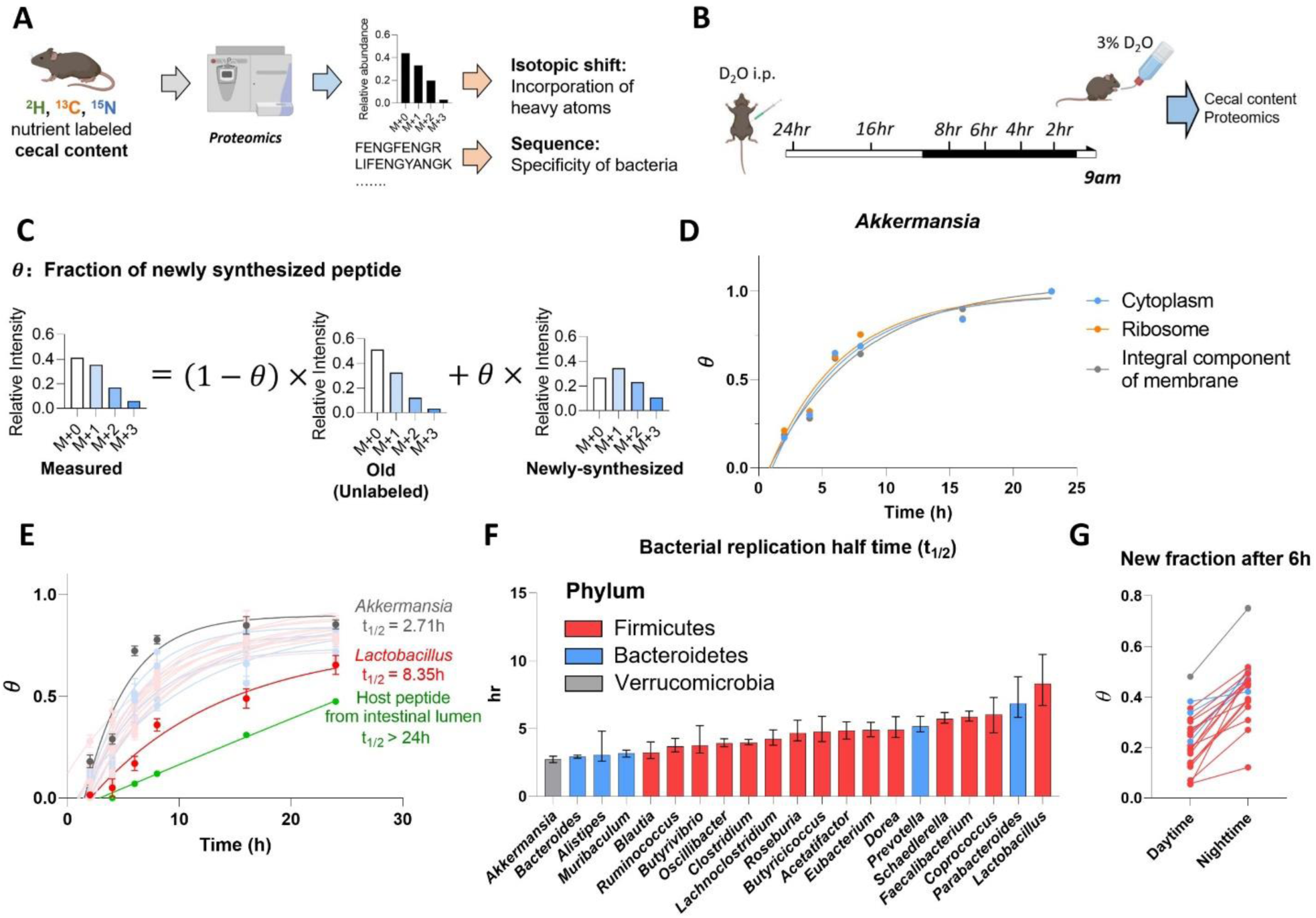
Growth rate of different gut bacterial genera quantified by isotope tracing. (A) Experimental approach for isotope tracing into specific gut bacteria. Only peptides that are specific to a particular bacterial genus were examined. (B) Growth rate quantification using D_2_O. Mice received D_2_O by i.p. injection followed by D_2_O drinking water and cecal content labeling was measured over time by proteomics and metabolomics. (C) Calculation of newly synthesized peptide fraction (*θ*). The experimentally observed peptide mass isotope distribution was fit to a linear combination of unlabeled peptide (“old,” heavy forms from natural isotope abundance) and newly synthesized peptide (“new,” heavy forms from isotope labeling pattern of free cecal amino acids and from natural isotope abundance). (D) Different cellular compartments from the same bacterial genus show similar labeling rate. (E) Genus-specific growth rates were determined by a single exponential fitting, as a function of time, of θ (mean across both different peptides measured from that genus and replicate mice). Mean±s.e. N=5 mice for each time point. (F) Bacterial replication half time of different gut bacteria. Data are exponential fits ±s.e. (G) The gut bacteria synthesize protein in sync with the physiological feeding patterns of the host. The figure shows the average newly synthesized peptide fraction (*θ*) for different gut bacterial genera after D_2_O labeling during daytime vs nighttime. Each line connects the daytime and nighttime measurements for one genus. Mean, N = 5 mice for daytime and for nighttime.

To quantify protein synthesis in different gut microbial genera, we used D_2_O tracing (Holmes et al., 2015; O’Brien et al., 2020). To achieve steady-state labeling of body water, we gave mice D_2_O by bolus injection followed by mixing it into drinking water. Peptide labeling in the cecal contents was then measured by proteomics (**Figure 4B**).

A key technical challenge in using proteomics to read out metabolic activity is the complexity, arising from natural isotope abundances, of peptide mass spectra. We used liquid chromatography-high resolution mass spectrometry to obtain the full scan (MS1) mass isotope distribution for each peptide of interest, with MS/MS analysis of the unlabeled form used to determine the peptide’s identity. We then calculated, based on the mass isotope distribution, the fraction of peptide that was newly synthesized (*θ*). To this end, first, we calculated the mass isotope distribution of unlabeled peptides based on natural isotope abundances (“old”). Second, we calculated the expected mass isotope distribution of a newly-synthesized peptide generated from cecal free amino acids, whose labeling we experimentally measured by metabolomics. Then, we determined the fraction of newly synthesized (*θ*) by linear interpolation between the “old” and “newly synthesized” spectra (**Figure 4C**).

For each bacterial genus, we measured the newly synthesized fraction (*θ*) for a minimum of 5 peptides, with abundant gut bacteria yielding θ for over 100 characteristic peptides. Irrespective of their intracellular location, different peptides from the same bacterial genus tended to label at a similar rate, suggesting that the peptide labeling rate largely reflects bacterial growth rate (**Figure 4D, S5A**). Labeling rate varied across bacterial genera, with a half doubling time ranging from 2.5 h for *Akkermansia* to 8 h for *Lactobacillus*, which still markedly exceeded the labeling rate of host intestinal proteins (> 24 h half doubling time) (**Figure 4E-F, Figure S5B**).

Our prior analyses revealed that the microbiome is fed substantially by dietary components. Accordingly, we hypothesized that microbial growth synchronizes with physiological feeding, which in mice occurs mainly during the nighttime. To assess the diurnal rhythm of gut bacterial protein synthesis, mice were given D_2_O for 6 h intervals throughout the diurnal cycle, followed by proteomic analysis of their cecal contents. Every measured bacterial genus showed greater protein synthesis during nighttime than daytime (**Figure 4G**). Thus, gut bacteria synthesize protein in sync with the physiological feeding patterns of the host.

### Preferred carbon sources differ across gut bacteria

Next, we quantitated the carbon feedstocks of different microbes, by combining ^13^C-nutrient labeling and proteomics. Each ^13^C-labeled nutrient (dietary inulin, dietary algal protein, or circulating lactate) was provided for 24 hours, which is sufficient to achieve steady-state labeling in the gut bacteria. Our analysis strategy involved two steps: first, we calculated, based on each genus-specific peptide’s observed mass isotope distribution, its relative ^13^C-enrichment (γ) compared to that of cecal free amino acids (**Figure 5A**). Mathematically, this calculation is identical to the calculation of θ in the D_2_O case, except here, the tracer is a particular ^13^C-labeled nutrient, which unlike D_2_O is used preferentially by certain bacterial genera. The observed peptide’s relative ^13^C-enrichment multiplied by the average contribution of that ^13^C-tracer to the gut microbial amino acids pool (*L*_*AA_avg*←*nutrient*_) gives a quantitative measure of the tracer’s contribution to the observed genus-specific peptide. Averaging across such peptides gives a fractional contribution of the ^13^C-labeled nutrient to protein synthesis in a bacterial genus.

**Figure 5.**
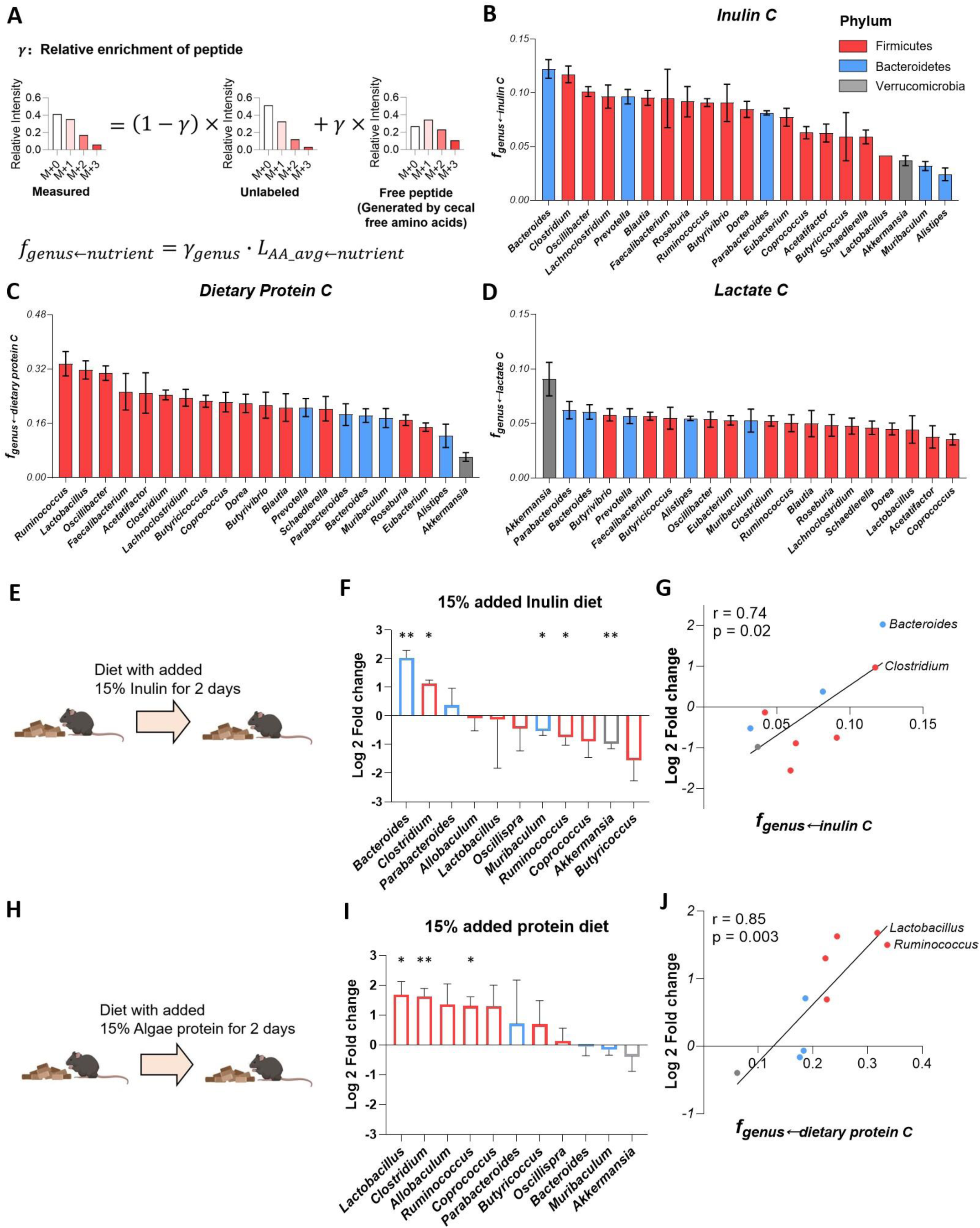
Preferred carbon sources differ across gut bacteria. (A) Calculation of peptide relative ^13^C-enrichment (*γ*) and carbon contribution from the tracer to a bacterial genus (*f*_*genus←nutrient*_). First, the experimentally observed peptide mass isotope distribution was fit to a linear combination of unlabeled peptide (heavy forms from natural isotope abundance) and a peptide made from free cecal amino acids (heavy forms from isotope labeling pattern of free cecal amino acids and from natural isotope abundance), yielding γ. Then, *f*_*genus←nutrient*_ was determined by correcting for the fractional contribution of that tracer to the cecal free amino acid pools. (B) Carbon contribution of dietary inulin across bacterial genera. Mean±s.e. N=4 mice. (C) Carbon contribution of dietary algal protein across bacterial genera. Mean±s.e. N=6 mice. (D) Carbon contribution of circulating lactate across bacterial genera. Mean±s.e. N=7 mice. (E) Experimental scheme of high-inulin diet feeding followed by 16S ribosomal RNA sequencing. (F) Genus-level microbiota composition changes after high-inulin diet. The genera increased after high-inulin diet prefer inulin in (B). Mean±s.e. N=3 mice. *P<0.05 and **P<0.01 by two-sided Student’s t-test. (G) Correlation between genera abundance changes and carbon-source preference. (H-J) As in (E - G), for algal protein-supplemented diet.

Using this method, we measured feedstocks of the bacterial genera that were detected in every proteomics experiment. We observed marked differences in nutrient preferences across members of the microbiota. For example, *Bacteroides* and *Clostridium* use over four-fold more inulin than *Akkermansia, Muribaculum*, or *Alistipes* (**Figure 5B**). Overall, bacteria from the phylum Firmicutes, used more dietary protein than did Bacteroidetes (Firmicutes 0.237 ± 0.052; Bacteroidetes 0.175 ± 0.031, p = 0.02).

*Akkermansia*, which is generally considered a health-promoting gut microbe, used among the least dietary inulin and protein (**Figure 5B-C**). In contrast, it used by far the most circulating lactate from the host (**Figure 5D**).

We were curious whether these bacterial nutrient preferences predict microbiome composition changes upon dietary changes. To explore this possibility, we fed mice an inulin-enriched or algal protein-enriched diet for two days and measured microbiome composition by 16S rRNA sequencing. Major bacteria genera with a relative abundance > 0.5% in 16S rRNA sequencing were examined. *Bacteroides*, the top consumer of ^13^C-inulin, increased by 4-fold after high inulin diet (**Figure 5E-G**). *Clostridium*, another high inulin consumer, also increased by 2-fold. Other genera that use less inulin carbon were either unchanged or slightly decreased. Similar consistency between microbes’ nutrient preference and relative abundance changes was observed in mice fed the algal protein-enriched diet (**Figure 5H-J**). Carbon-source preference measured by proteomics (*f*_*genus*←*nutrient*_) correlates with abundance change following a diet shift measured by 16S sequencing, for both the inulin and algal protein conditions (**Figure 5G, J**). Thus, the nutrient preferences of different gut bacteria help explain microbiome compositional changes following dietary manipulations (David et al., 2014).

### Firmicutes consume dietary protein while Bacteroidetes consume secreted host protein

Lastly, we turned to the nitrogen source preferences of different gut bacteria, comparing ^15^N-labeled dietary protein feeding to ^15^N-urea infusion. The analytical approach was identical to that employed above for carbon source preferences. Bacterial genera that highly use carbon from dietary protein also highly use nitrogen from dietary protein, consistent with amino acids from dietary protein being assimilated intact in bacterial proteomes **(Figure 6A, S6A)**.

**Figure 6.**
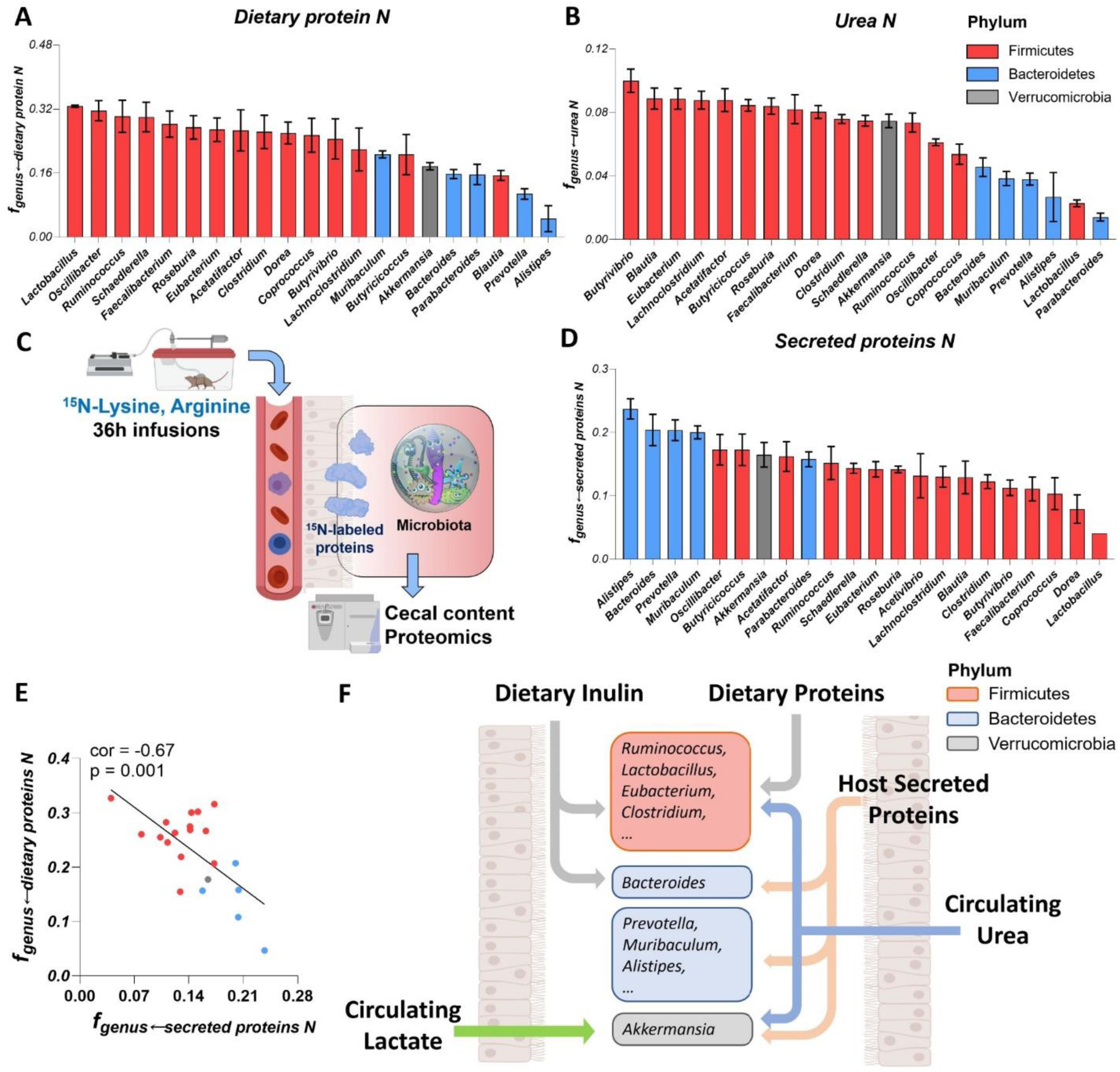
Firmicutes favor dietary protein while Bacteroidetes prefer secreted host protein. (A) Nitrogen contribution of dietary algal protein across bacterial genera. Mean±s.e. N=6 mice. (B) Nitrogen contribution of circulating urea across bacterial genera. Mean±s.e. N=6 mice. (C) Experimental schematic of long-term ^15^N-lysine and ^15^N-arginine infusion to probe the contribution of secreted host proteins to different bacterial genera. (D) Nitrogen contribution of secreted host proteins across bacterial genera. Mean±s.e. N=5 mice. (E) Negative correlation between *f*_*genus*←*secreted proteins N*_ and *f*_*genus*←*dietary proteins N*_. (F) Summary of carbon and nitrogen inputs to different gut bacteria. Firmicutes prefer dietary carbon sources (fiber and protein) and nitrogen from host circulating urea. Bacteroidetes heavily use dietary fiber, while using on host secreted proteins for nitrogen. Verrucomicrobia prefers host secreted nutrients, both protein and circulating small molecules (lactate, urea).

Conversely, among members of the phylum Firmicutes, genera preferring urea nitrogen tended to be avid inulin users, i.e. to synthesize their own amino acids using inulin and urea **(Figure 6B, S6B)**. Finally, again among Firmicutes, we also saw the expected trade-off where some genera prefer nitrogen from dietary protein, and others from circulating urea **(Figure S6C)**. The most intriguing observation, however, was that bacteria from the phylum Firmicutes used more nitrogen both from dietary proteins and from circulating urea than did Bacteroidetes (Dietary proteins: Firmicutes 0.263 ± 0.044; Bacteriodetes 0.135 ± 0.006, p = 7.7 × 10^−5^ ; Urea: Firmicutes 0.076 ± 0.019; Bacteriodetes 0.033 ± 0.012, p = 0.0001) (**Figure 6A-B**).

The low use of both dietary protein and circulating urea nitrogen by Bacteroidetes raised a key question: How do Bacteroidetes get nitrogen? It has been shown that some members of gut microbiome (e.g. *Bacteroides* and *Akkermansia*) are capable of digesting host secreted proteins such as mucins (Berry et al., 2013; Reese et al., 2018). We hypothesized that host secreted proteins are a key source of Bacteroidetes nitrogen. To probe this possibility, we performed long-term (36 h) ^15^N-labeled lysine and arginine infusions to label host proteins in the colon (**Figure 6C and Figure S7A-D**). Lysine and arginine do not directly feed the microbiome (**Figure 1E**) but did make a discernible contribution after the long-term infusion, consistent with the labeling occurring via host proteins. Such labeling occurred preferentially in Bacteriodetes peptides (**Figure 6D**). *Akkermansia*, consistent with its mucin degrading capability, is another user of host secreted proteins. The nitrogen contributions from dietary and secreted host proteins were anti-correlated, consistent with some gut bacteria preferentially consuming dietary protein, and others host protein (**Figure 6E**). Interestingly, bacterial genera with a higher preference for dietary protein, which is dependent on host feeding, tend to grow more differently between daytime and nightime, while genera that prefer host secreted proteins, grow at a similar rate throughout a day. (**Figure S6D-E**). Thus, dietary proteins and circulating urea are the major nitrogen feedstock of Firmicutes, while secreted host proteins provide nitrogen to Bacteroidetes.

## Discussion

As for most microbial communities, the composition of the gut microbiome is shaped by nutrient availability. Here we developed quantitative isotope tracing approaches to measure the nutrient preferences of gut bacteria. In addition to dietary fiber and secreted host proteins, we establish dietary protein and circulating host lactate, 3-hydroxybutyrate, and urea as important nutrients feeding gut bacteria. Importantly, we rule out direct contributions from other circulating host nutrients, like glucose and amino acids, to the colonic microbiome.

A key technical achievement is enabling tracing from different carbon and nitrogen sources into bacteria-specific peptides, thereby revealing the nutrient preferences of different bacteria within the complex and competitive gut lumen environment. We find that Firmicutes and Bacteroidetes differ systematically in their utilization of host secreted protein versus dietary protein: Firmicutes tend to acquire amino acids from dietary protein, while Bacteroidetes rely more on secreted host protein (**Figure 6F**). This may relate to different localization of bacteria within the colon, either in terms of central versus peripheral (closer to host mucus) or distal versus proximal (closer to incoming food remnants) (Albenberg et al., 2014; Li et al., 2015; Yasuda et al., 2015).

Within these two major families of gut bacteria, we found marked disparities in the use of dietary fiber as a carbon source. The most abundant Bacteroidetes’ genus is *Bacteroides*, and it was the most avid assimilator of fiber (inulin). In contrast, other types of bacteria in the same phylum hardly consumed inulin. Likewise, some Firmicutes like *Clostridium* avidly used fiber, while others did not. Strikingly, feeding a fiber-enriched diet led to an increased abundance of *Bacteroides* and *Clostridium*, the precise genera that most actively assimilate fiber based on isotope tracing.

A similar trend was observed in the case of dietary supplementation with algal protein: Firmicutes, which actively use such protein, tended to increase in abundance. Algal protein (the only type commercially available in bulk in ^13^C-labeled form) may be particularly hard for mammals to digest. This is reflected in the limited appearance of ^13^C-labeled amino acids from algal protein in the portal circulation, and instead extensive passage from the intestine into the colon. This influx of dietary protein to the microbiome was a major contributor to secreted microbiome metabolites: dietary algal protein provided a portion of the carbon in the most abundant microbiome products (e.g., SCFAs) and was the main source of most other microbiome-derived metabolites. An important future question is whether the nature of dietary protein (e.g. plant or animal-based) impacts passage through the small intestine to the colonic microbiome and thereby shapes microbiome composition or metabolite secretion (Madsen et al., 2017; Wali et al., 2021).

Host circulating metabolite levels may also impact microbiome nutrient access and ultimately composition. Here we show such effects are likely limited to the few host metabolites that meaningfully penetrate the microbiome: urea, 3-hydroxybutyrate, and lactate. Among them, lactate was recently shown to feed the gut microbiome in human marathon runners (Scheiman et al., 2019). Among gut bacteria, *Akkermansia* most avidly use circulating lactate. *Akkermansia* are mucin degraders, and their proximity to the gut epithelial wall may augment their access to lactate from the host circulation. *Akkermansia* are more abundant in athletes, and exercise increases their levels in mice and human (Liu et al., 2017; Munukka et al., 2018). A possible mechanism involves increased circulating lactate levels following exercise directly feeding *Akkermansia*. Whether lactate-induced *Akkermansia* growth in part mediates beneficial effects of exercise is an important open question.

Ultimately, manipulating the microbiome requires understanding which nutrients different bacteria consume, and how such consumption impacts microbiome composition and product secretion. Through isotope tracing, including proteomic measurements that offer bacterial genus specificity, we provide foundational knowledge about which nutrients feed the gut microbiome, and which bacteria prefer which nutrients. Currently, our measurements are limited to young mice, fed typical chow with or without fiber or algal protein supplementation. Furthermore, they are limited to the genus level and to genera that we detect by proteomics and/or sequencing. Nevertheless, we cover many important gut microbiome genera. Moreoever, the methodologies developed here are poised for broader application, to eventually contribute to the holistic and quantitative understanding of the diet-microbiome-health connection.

## Methods

### Mouse gavage and labeled nutrient feeding

Mouse studies followed protocols approved by the Princeton University Animal Care and Use Committee. Unless otherwise indicated, mice were group-housed on a normal light-dark cycle (8:00-20:00) with free access to water and chow. For the ^13^C-nutrient gavage experiments, 7-9-week-old male C57BL/6NCrl mice (strain 027; Charles River Laboratories) were fasted at 9 am and received a 1:2:4 mixture of inulin, protein/amino acids, and starch (0.5 g kg^-1^ inulin, 1 g kg^-1^ protein/amino acids 2g kg^-1^ starch dissolved in water) at 3 pm via oral gavage with a plastic feeding tube (Instech Laboratories). Food was given back at 8 pm.

For the experiments involving labeled nutrient feeding, the labeled diet was prepared by adding ^13^C/^15^N-nutrients to a diet mixture premix (modified from normal diet with reduced protein, inulin, and starch content, Research diets Inc, D20030303). The final enrichment for each labeled dietary nutrient was 10% - 25% (with observed labeling corrected by dividing by the fraction dietary nutrient labeled). The contribution of each dietary nutrient to metabolites is calculated by the metabolite labeling enrichement normalized to the final enrichment of each labeled dietary nutrient. All diets shared the same final macronutrient composition (40% starch, 20% protein or amino acids, 7.5% inulin and 2.5% cellulose). Male C57BL/6NCrl mice (7-9-week-old, strain 027, Charles River Laboratories) were first adapted to a non-labeled diet (of identical composition to the subsequent labeled diet) for 10 days, and then fed labeled diet for 24 h prior to sacrifice.

For the deuterium water drinking experiment, mice were administered a bolus intraperitoneal injected of D_2_O (1.26 % w/w relative to body weight), followed by having ad lib access to 3% D_2_O drinking water.

### Intravenous infusions

To quantify contribution of circulating nutrients to microbiota metabolism, 9-11-week-old C57BL/6 mice were catheterized in house in the right jugular vein. The mice were infused with carbon or nitrogen-labeled tracer starting at 3:30 pm without any fasting. Infusion rate was 0.1 ul/min/g. Infusion solutions are described in **Table S3**. Overnight (24 h) infusions both started and finished around 9 am. The contribution of circulating nutrient to each metabolite is calculated by the metabolite labeling enrichment normalized to the average tracer serum enrichment throughout 24 hr.

### Antibiotics treatment

To deplete the mouse resident microbiome, an antibiotic drinking water protocol was used. In brief, mice were treated with a cocktail of antibiotics (1 g/L ampicillin, 1 g/L neomycin, 1 g/L metronidazole, and 1 g/L vancomycin) in both their drinking water 14 days. To make the drinking water more palatable, 5% aspartame was added The effectiveness of antibiotics treatments were verified by observing much lower SCFAs in the feces by LC-MS.

### Sample collection

Systemic blood samples (∼6 µl) were collected by tail bleeding. For sampling from tissue-specific draining veins, a mouse was put under anesthesia and different tissue veins were exposed, and blood samples were pulled with an insulin syringe (BD insulin syringes, # SY8290328291) insertion into the vein. Successful isolation of portal vein was confirmed by much higher (> 10x) concentrations of SCFAs and secondary bile acids (deoxycholic acid and lithocholic acid) than systemic vein; hepatic vein was confirmed by much lower secondary bile acids, SCFAs and higher glucose, 3-hydroxybutyrate levels compared to portal vein. Mouse urine was collected from the urinary bladder using a syringe. All serum samples were placed on ice without anticoagulant for 15 min, and centrifuged at 16,000 x g for 15 min at 4 C.

Tissues were harvested by quick dissection and snap freezing (<5 sec) in liquid nitrogen with a pre-cooled Wollenberger clamp; intestinal contents were removed before clamping. For cecal content sampling, the mouse cecum was first removed and cut on the surface, then the cecal content was sequeezed out using a tweezer followed by freeze clamping. Whole liver, intestine, and intestinal contents were collected and grounded to homogenous powder. To sample fresh feces, the mouse belly was gently massaged to induce defecation and fresh feces were freeze clamped. For long-term feces collection, a mouse was transferred to a new cage and mouse fecal pellets on the bedding were collected every 1∼2 h and freeze clamped. Serum, tissue, and feces samples were kept at −80 °C until further analysis.

### 16S rRNA gene amplicon sequencing and analysis

Extraction of Bacterial DNA from cecal or fecal samples was performedusing the Power Soil DNA Isolation kit (QIAGEN). A section of the 16S rRNA gene (∼250 bp, V4 region) was amplified, and Illumina sequencing libraries were prepared from these amplicons according to a previously published protocol and primers (Caporaso et al., 2012). Libraries were further pooled together at equal molar ratios and sequenced on an Illumina HiSeq 2500 Rapid Flowcell or MiSeq as paired-end reads. These reads were 2×150 bp with an average depth of ∼20,000 reads. Also included were 8 bp index reads, following the manufacturer’s protocol (Illumina, USA). Pass-Filter reads were generated from raw sequencing reads using Illumina HiSeq Control Software. Samples were de-multiplexed using the index reads. The DADA2 plugin within QIIME2 version 2018.6 was used to inferred Amplicon sequencing variants (ASVs) from the unmerged paired-end sequences (Bolyen et al., 2019; Callahan et al., 2016). The forward reads were trimmed at 150 bp and the reverse reads trimmed at 140 bp, with all other DADA2 as default. Taxonomy was assigned to the resulting ASVs with a naïve Bayes classifier trained on the Greengenes database version 13.8, with only the target region of the 16S rRNA gene used to train the classifier (Bokulich et al., 2018; McDonald et al., 2012). Downstream analyses were performed MATLAB (Hunter, 2007; McKinney, 2010).

### Metabolite extraction

For serum samples, 3 ul serum was added to 90 ul methanol and incubated on ice for 10 min, followed by centrifugation at 17,000 × g for 10 min at 4°C. The supernatant was transferred to an MS vial until further analysis. For tissues and feces samples, frozen samples were first ground at liquid nitrogen temperature with a cryomill (Restch, Newtown, PA). The resulting tissue powder was extracted with 40:40:20 methanol: acetonitrile: water (40 ul extraction solvent per 1 mg tissue) for 10 min on ice, followed by centrifugation at 17,000 x g for 10 min, the supernatant was transferred to a MS vial until further analysis.

### Measurements of metabolites, protein, and polysaccharides

To measure metabolites in serum, tissue and feces samples, a quadrupole orbitrap mass spectrometer (Q Exactive; Thermo Fisher Scientific) was coupled to a Vanquish UHPLC system (Thermo Fisher Scientific) with electrospray ionization and scan range m/z from 60 to 1000 at 1 Hz, with a 140,000 resolution. LC separation was performed on an XBridge BEH Amide column (2.1×150 mm, 2.5 μm particle size, 130 Å pore size; Waters Corporation) using a gradient of solvent A (95:5 water: acetonitrile with 20 mM of ammonium acetate and 20 mM of ammonium hydroxide, pH 9.45) and solvent B (acetonitrile). Flow rate was 150 μl/min. The LC gradient was: 0 min, 85% B; 2 min, 85% B; 3 min, 80% B; 5 min, 80% B; 6 min, 75% B; 7 min, 75% B; 8 min, 70% B; 9 min, 70% B; 10 min, 50% B; 12 min, 50% B; 13 min, 25% B; 16 min, 25% B; 18 min, 0% B; 23 min, 0% B; 24 min, 85% B; and 30 min, 85% B. Injection volume was 5-10 μl and autosampler temperature was set at 4°C. For cysteine measurement, samples were derivatized before measurement as follows: Serum, cecal content or feces samples were extracted and centrifuged. To the supernatant, 2 mM N-ethylmaleimide was added and incubated at room temperature for 20 min. The resulting mixture was transferred to a MS vial. Derivatized cysteine has a m/z at 245.06015 in negative mode.

To quantify the metabolite concentration in serum and tissue samples, either isotope spike-in or standard spike-in was performed. For isotope spike-in, known concentrations of isotope-labeled standard were added to the serum or tissues extraction solution, then the concentration was calculated by the ratio of labeled and unlabeled metabolites. When isotope standard is not available, a serially diluted non-labeled standard was added, and a linear fitting between measured total ion count and added concentration of standard was generated. Then, the concentration of endogenous metabolite was determined by the x intercept of the fitting line.

Starch and inulin were measured by acid hydrolysis and LC-MS. In brief, 5-10 mg sample was mixed with 10 ul 2 M hydrochloric acid, and samples were incubated at 80°C for 2 h. After cooling down, the resulting mixture was neutralized with 12 μl saturated sodium bicarbonate, followed with 88 μl 1:1 acetonitrile: methanol solution. After centrifugation at 17,000 × g for 10 min at 4°C, the supernatant was transferred to a MS vial. Inulin and starch concentration in samples was inferred from total ion count of fructose and glucose, respectively.

SCFAs and BCFAs were derivatized and measured by LC-MS. Serum (5 μl) or tissue samples (∼10 mg) were added to 100 μl derivatizing reagents containing 12 mM 1-Ethyl-3-(3-dimethylaminopropyl) carbodiimide, 15 mM 3-Nitrophenylhydrazine hydrochloride acid and pyridine (2% v/v) in methanol. The reaction was incubated at 4°C for 1 h. Then, the reaction mixture was centrifuged at 17,000 g for 10 min. 20 μl supernatant was quenched with 200 μl 0.5 mM beta-mercaptoethanol in 0.1% formic acid water. After centrifugation at 17,000 g for 10 min, the supernatant was transferred to MS vials until further analysis. The measurement of SCFAs and BCFAs are performed using the same Q Exactive PLUS hybrid quadrupole-orbitrap mass spectrometer with different column and LC setup. LC separation was on Acquity UPLC BEH C18 column (2.1 mm x 100 mm, 1.7 5 μm particle size, 130 Å pore size, Waters, Milford, MA) using a gradient of solvent A (water) and solvent B (methanol). Flow rate was 200 μL/min. The LC gradient was : 0 min, 10% B; 1 min, 10% B; 5 min, 30% B; 11 min 100% B; 14 min, 100% B; 14.5 min 10% B; 22 min 10 % B. Autosampler temperature was 5 °C, column temperature was 60 °C and injection volume was 10 μl. Ion masses for derivatized acetate, propionate, butyrate, iso-butyrate, valeric acid, isovaleric acid, 2-methylbutyrate, 4-methylvaleric acid were 194.0571, 208.0728, 222.0884, 222.0884, 236.1041, 236.1041, 236.1041, 250.1197 in negative mode, respectively.

### Proteomics sample preparation

Proteomics samples were prepared mostly as previously described (Gupta et al., 2018; Wühr et al., 2014). Mouse cecal samples (10 mg each) were dissolved in 400 ul lysis buffer (6M guanidium chloride, 2% cetrimonium bromide, 5 mM dithiothreitol, 50 mM (4-(2-hydroxyethyl)-1-piperazineethanesulfonic acid) (HEPES), pH 7.2). Then the sample mixture was put on ice and sonicated for 10 cycles (30 s on and 30 s off cycle, amplitude 50%) by a sonicator (Qsonica), followed by centrifugation at 20,000 × g for 20 min at 4 °C. The supernatant was taken and alkylated with 20 mM N-ethylmaleimide for 20 min at room temperature, 5 mM dithiothreitol was added to quench the excessive alkylating reagents. Proteins were purified by methanol-chloroform precipitation. The dried protein pellet was resuspended in 10 mM EPPS (N-(2-Hydroxyethyl) piperazine-N’-(3-propanesulfonic acid)) at pH 8.5 with 6 M guanidine hydrochloride. Samples were heated at 60°C for 15 min and protein concentration was determined by BCA assay (Pierce BCA Protein Assay Kit, Thermo Scientific). The protein mixture (30∼50 µg) was diluted with 10 mM EPPS pH 8.5 to 2 M GuaCl and digested with 10 ng/µL LysC (Wako) at room temperature overnight. Samples were further diluted to 0.5 M GuaCl with 10 M EPPS pH 8.5 and digested with an additional 10 ng/µL LysC and 20 ng/µL sequencing grade Trypsin (Promega) at 37°C for 16 h. Samples were desalted using a SepPak cartridges (Waters) and then vacuum-dried and resuspended in 1% formic acid before mass spectrometry analysis.

### Proteomics peptide measurement

Samples were analyzed on an EASY-nLC 1200 (Thermo Fisher Scientific) HPLC coupled to an Orbitrap Fusion Lumos mass spectrometer (Thermo Fisher Scientific) with Tune version 3.3. Peptides were separated on an Aurora Series emitter column (25 cm × 75 μm ID, 1.6 μm C18) (Ionopticks, Australia) and held at 60°C during separation using an in-house built column oven over 180 min, applying nonlinear acetonitrile gradients at a constant flow rate of 350 nL/min. The Fusion Lumos was operated in data dependent mode. The survey scan was performed at a resolution setting of 120k in orbitrap, followed by MS2 duty cycle of 1.5 s. The normalized collision energy for CID MS2 experiments was set to 30%.

Solvent A consisted of 2% DMSO (LC-MS-grade, Life Technologies), 0.125% formic acid (98%+, TCI America) in water (LC-MS-grade, OmniSolv, VWR), solvent B of 80% acetonitrile (LC-MS-grade, OmniSolv, Millipore Sigma), 2% DMSO and 0.125% formic acid in water. The following 120 min-gradient with percentage of solvent B were applied at a constant flow rate of 350 nL/min after thorough equilibration of the column to 0% B: 0% – 6% in 5 min; 6 – 25% in 160 min; 25% –100% in 10 min; 100% for 5 min. For electrospray ionization, 2.6 kV were applied between minutes 1 and 113 (or minutes 1 and 83 for fractionated samples) of the gradient through the column. To avoid carry-over of peptides, 2,2,2-trifluoroethanol (>99% Reagent plus, Millipore Sigma) was injected in a 30 min wash between each sample.

### Proteomics data analysis

The data was analyzed using GFY software licensed from Harvard (Nusinow et al., 2020). Thermo Fisher Scientific. raw files were converted to mzXML using ReAdW.exe. MS2 spectra assignment was performed using the SEQUEST algorithm v.28 (rev. 12) by searching the data against the combined reference proteomes for *Mus Musculus, Bos Taurus*, and all the abundant bacterial families detected in 16S rRNA sequencing (*Bacteroidaceae, Porphyromonadaceae, Prevotellaceae, Rikenellaceae, Muribaculaceae, Lachnospiraceae, Ruminococcaceae, Erysipelotrichaceae, Oscillospiraceae, Clostridiaceae, Eubacteriaceae, Lactobacillaceae* and *Verrucomicrobiaceae*) acquired from Uniprot on Jan 2021 (SwissProt + Trembl) along with common contaminants such as human keratins and trypsin. The target-decoy strategy was used to construct a second database of reverse sequences that were used to estimate the peptide false discovery rate (Elias and Gygi, 2007). A 20 ppm precursor ion tolerance with the requirement that both N- and C-terminal peptide ends are consistent with the protease specificities of LysC and Trypsin was used for SEQUEST searches, two missed cleavage was allowed. NEM was set as a static modification of cysteine residues (+125.047679 Da). An MS2 spectral assignment false discovery rate of 0.5% was achieved by applying the target decoy database search strategy. Linear Discriminant analysis was used for filtering with the following features: SEQUEST parameters XCorr and unique ΔXCorr, absolute peptide ion mass accuracy, peptide length and charge state. Forward peptides within three standard deviations of the theoretical m/z of the precursor were used as positive training set. All reverse peptides were used as negative training set. Linear Discriminant scores were used to sort peptides with at least seven residues and to filter with the desired cutoff. Furthermore, we performed a filtering step on the protein level by the “picked” protein FDR approach (Savitski et al., 2015). Protein redundancy was removed by assigning peptides to the minimal number of proteins which can explain all observed peptide, with above-described filtering criteria.

To quantify the intensities of all the isotopic peaks of the peptides, we used raw intensity. Missed cleavage peptides (more than one K or R in the peptide) and low signal to FT-noise peptides (M_0_ S/N < 20) were removed. Peptide phylogenetic assignment was performed using Unipept (Gurdeep Singh et al., 2019), ‘Equate I and L’ and ‘Advanced missed cleavage handling’ were not selected. Only peptides that are specific at a genus level were used for further analysis.

### Quantification of newly-synthesized fraction of peptide

To determine the newly synthesized fraction of a bacterial peptide in D_2_O drinking water experiment, we first measured the cecal content free amino acids deuterium labeling pattern using metabolomics. Then, for each peptide, we simulated the expected isotope envelope pattern if the peptide were old, i.e., unlabeled with deuterium (*I*_*old*_), versus if it were newly synthesized by taking up free amino acids from the cecal content (*I*_*new*_). *I*_*old*_ was calculated based on the peptide’s molecular formula and ^13^C, ^15^N, ^2^H, ^17^O, _18_O, ^32^S, ^33^S and ^36^S natural abundance. *I*_*new*_ was calculated based on the peptide’s sequence and experimentally observed labeling of the corresponding cecal free amino acids (after natural isotope correction), and the natural isotope abundance of the unlabeled atoms in the peptide’s formula. The simulation of expected peptide isotope distribution and fitting was performed using a MATLAB code: https://github.com/xxing9703/pepMID_simul. Exact mass isotopic peaks with appreciable abundances were bundled by nominal mass into fraction M+0, M+1, …M+n, constituting the final simulated spectrum. A least square fit was used to find the scalar θ that best fit the measured peptide isotopic distribution (*I*_*measured*_) to a linear combination of *I*_*old*_ and *I*_*new*_:

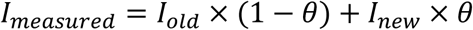

The root mean square error was determined for each peptide fitting, and any fitting with a root mean square error > 1% was removed. For genus-level turnover quantification, only genera with more than two measurements were kept in the analysis, with the median value across peptides reported.

### Quantification of contribution of labeled nutrient to peptide

To determine the contribution of a ^13^C-or ^15^N-labeled nutrient to a bacterial peptide, similar to the above approach, we first measured the cecal content free amino acids ^13^C-or ^15^N-labeling using metabolomics. Then, for each peptide, we simulated the expected isotope envelope pattern if the peptide were unlabeled (*I*_*unlabeled*_) versus if it were synthesized from free cecal amino acids (*I*_*free*_). A scalar γ (analogous to θ above) can then be determined by fitting the measured peptide isotope distribution (*I*_*measured*_) to a linear combination of *I*_*unlabeled*_ and *I*_*free*_. Note that γ will exceed 1 when a bacterial genus uses a particular nutrient in excess of that nutrient contribution’s to cecal free amino acids. Because the ^13^C- and ^15^N-labeling patterns are simpler than the D_2_O labeling patterns, in lieu of carrying out this fitting, we instead determined γ (with the same conceptual and mathematical meaning) using simple algebraic equations.

Specifically, we measured γ for each peptide as follows:

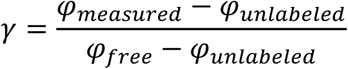

where (with the exception of ^13^C-protein feeding data, discussed immediately below) *φ* is the average number of extra neutrons in a given peptide (or simulated peptide), relative to the M+0 form. This was calculated based on the experimentally observed (or simulated, as above) fraction of M+0, M+1, M+2, and M+3, which account for > 90% of the isotopes for each peptide (with more heavily labeled forms too low abundance and noisy to contribute productively to the measurements):

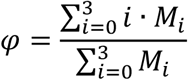

For the ^13^C-protein feeding experiments, the most readily detected labeled forms involve incorporation of a single midsized U-^13^C-amino acid, which manifests as M+5 or M+6 peptide labeling. Other isotopic forms were sufficiently noisier, as to render their inclusion unhelpful. Accordingly, we calculated γ based on *φ*^*′*^:

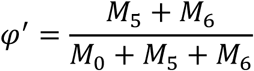

The above equations give nearly identical values for γ as fitting (as done to determine *θ*).

For genus-level measurements of feedstock contributions, only genera with more than 3 peptides measured per mousewas kept in the analysis, with the median value across peptides reported as

*γ*_*genus*_. Only genera that were consistently detected in proteomics, and the family of that genera detected (>0.5%) in 16S rRNA sequencing were analyzed. The product of *γ*_*genus*_ and the contribution of each nutrient to cecal free amino acids (*L*_*AA_avg*←*nutrient*_) was used to determine the contribution of each nutrient to bacterial genus (*f*_*genus*←*nutrient*_):

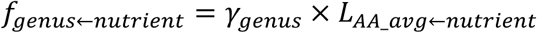

where the contribution of each nutrient to bacterial protein pool (*L*_*AA_avg*←*Nutrient*_) was calculated as the average labeling across amino acids, weighted based on their abundance in that genus’ protein and corrected for fraction of the nutrient interest labeled (*T*):

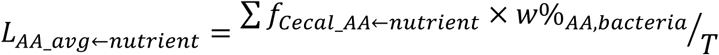

with *w*%_*AA,bacteria*_ taken from literature (Purser and Buechler, 1966).

### Statistical analysis

A two-tailed, unpaired student’s t-test was used to calculate P values, with P<0.05 used to determine statistical significance.

## Author contributions and information

X.Z., and J.D.R came up with general approach and X.Z. performed most of the experiments and data analysis. C.J. worked intensively with X.Z. to develop the experimental strategy. M.W. designed and enabled the proteomic measurements. X.X. wrote the MATLAB code. M.G., F.C.K., and M.D.N. contributed to proteomics method development. J.G.L. and M.S.D. provided microbiome expertise and performed 16S rRNA sequencing. A.R. assisted with isotope tracing. W.L. performed ammonia measurement. X.Z., C.J., and J.D.R. wrote the paper. All of the authors discussed the results and commented on the paper.

---

**Figure S1.**
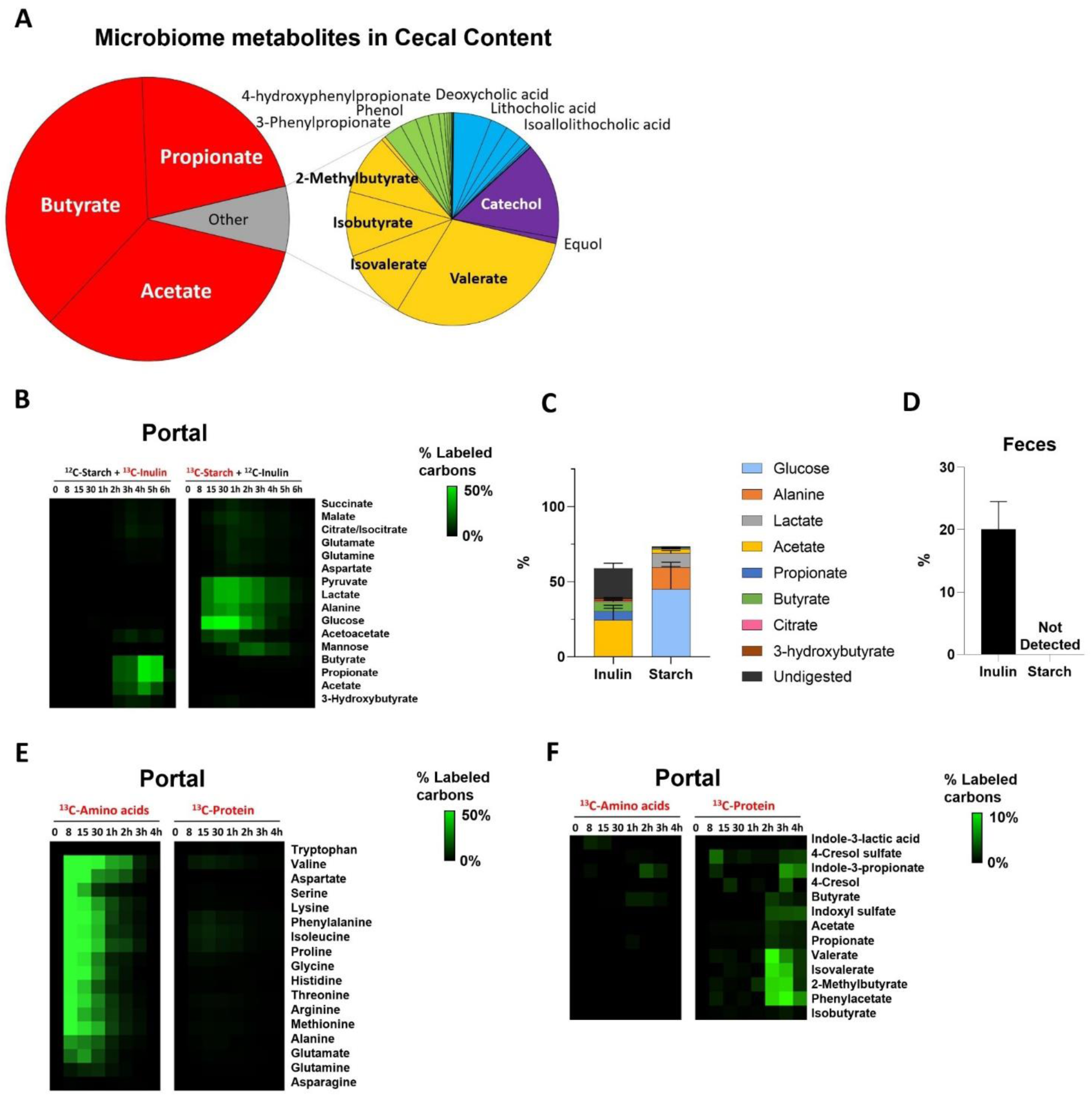
Microbiome consumes dietary fiber and protein. (A) Composition of the measured cecal microbial metabolome. The pie charts show the molar abundance of different gut microbiota-associated metabolites (N = 6 mice). (B) Heatmap showing the percentage of labeled carbon atoms in the indicated metabolites in portal circulation, following gavage of 4:2:1 starch: protein (or free amino acids): inulin, with the indicated nutrient labeled. Each data point is median of N = 3 mice. (C) Metabolic fate of inulin and starch. Stacked bars show the fraction of gavaged inulin and starch that is converted into each of the indicated metabolic products, with the undigested fraction being excreted in the feces (mean ± s.e., N = 3 mice). (E) Undigested carbohydrate in the feces. Graph shows the fraction of labeled hexose after cecal content hydrolysis, 12 h following gavage as above (mean ± s.e., N = 3 mice). (F) Heatmap showing the percentage of labeled carbon atoms in the indicated amino acids in portal circulation, following gavage of 4:2:1 starch: protein (or free amino acids): inulin, with the indicated nutrient labeled. Each data point is median of N = 3 mice. (G) As in (E), for microbiota-associated metabolites.

**Figure S2.**
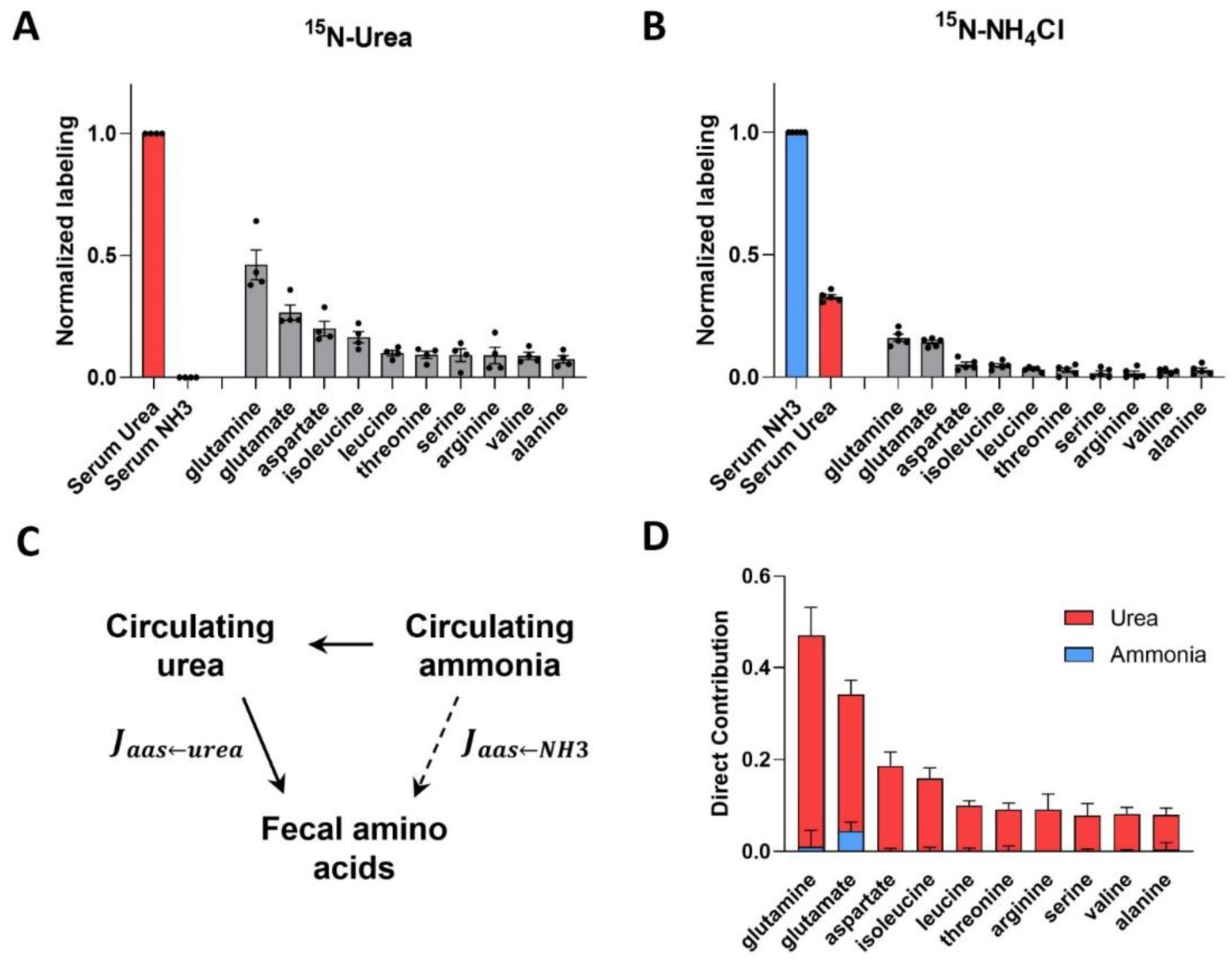
Circulating ammonia contributes to microbiota metabolism via circulating urea. (A) Normalized labeling of serum urea, ammonia and cecal amino acids after ^15^N-urea infusion (mean± s.e., N = 4 mice). Feces labeling fraction is normalized to serum infused tracer (urea) labeling fraction. (B) As in (A), for ^15^N-ammonia infusion (mean± s.e., N = 5 mice), (C) Model of direct contribution calculation from circulating urea and ammonia to fecal amino acids. (D) Direct contributions of circulating urea and ammonia to cecal amino acids (mean± s.e., N = 4 mice).

**Figure S3.**
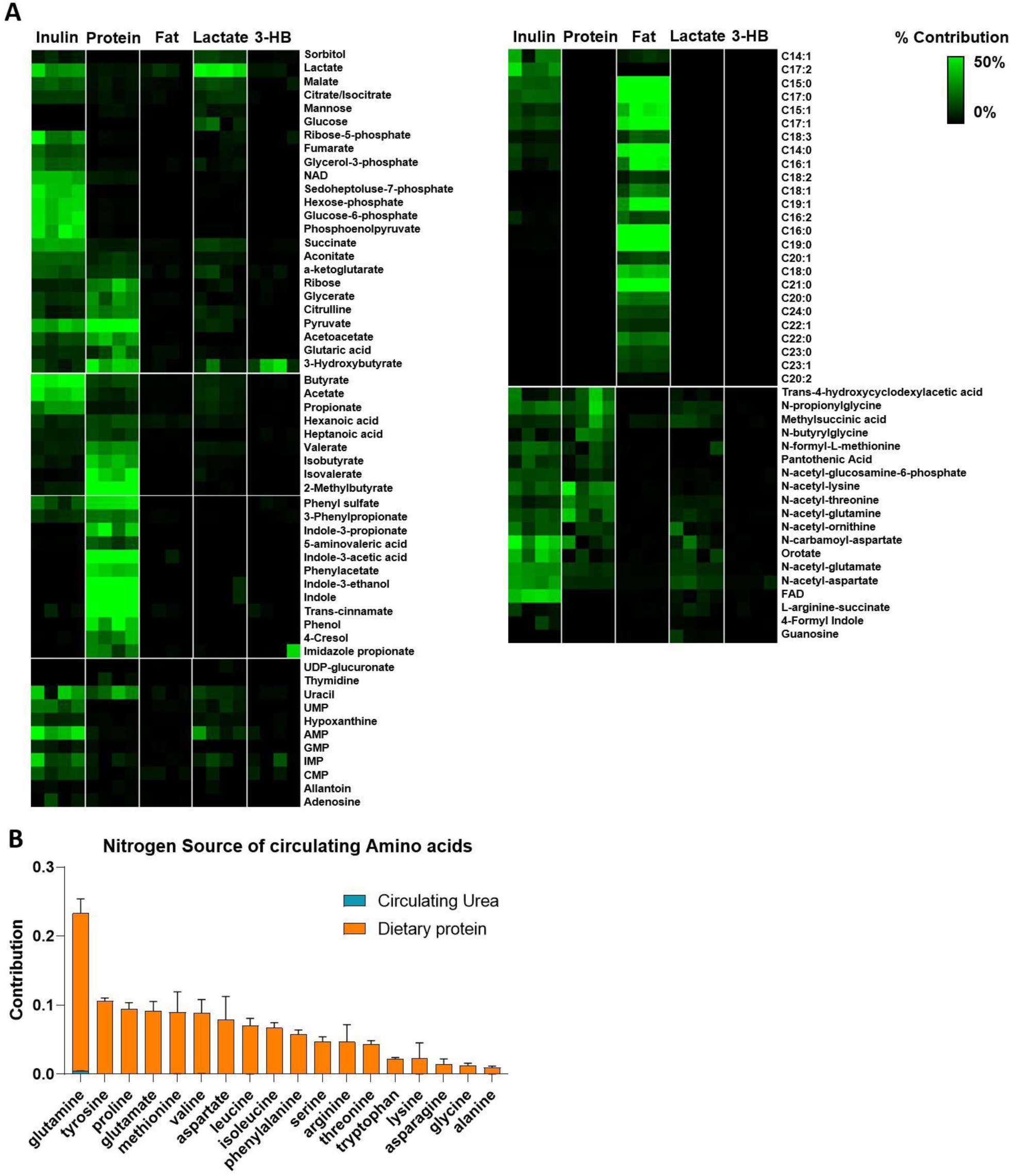
Quantitative analysis of dietary and circulating nutrient contributions to gut microbiome metabolism. (A) Heat maps showing the contribution of dietary or circulating nutrients to cecal metabolites. For experimental design, see Figure 3. N = 4 mice. (B) Amino acids synthesized in the gut microbiome, stay in the microbiome, as urea contributes to microbiome amino acids but not host circulating amino acids. Mean ± s.e. N = 4 mice.

**Figure S4.**
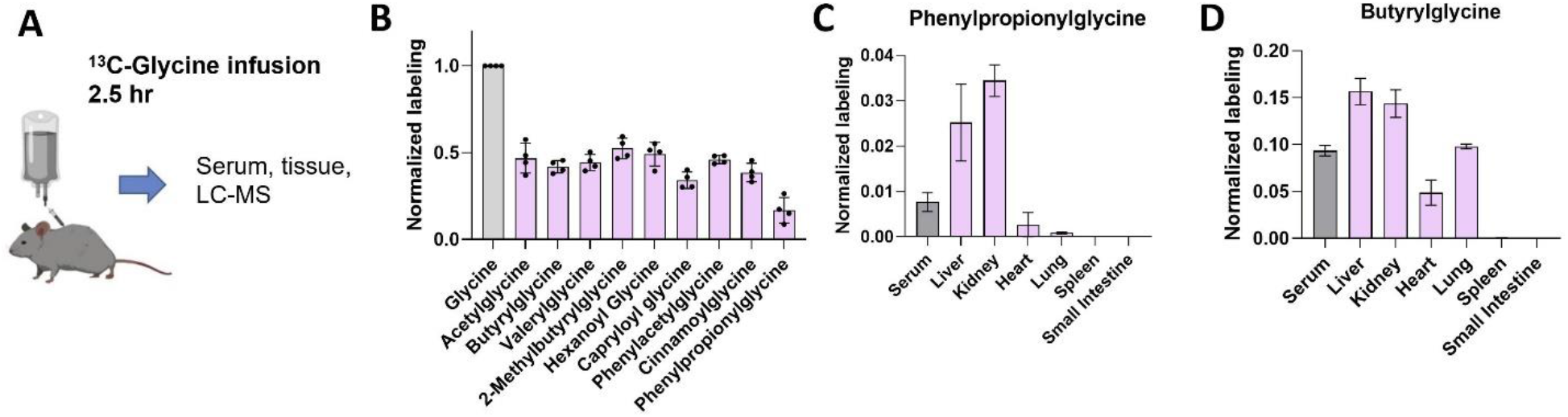
Liver and kidney use circulating glycine to synthesize acyl-glycines. (A) Experimental design. Mice were intravenously infused with [U-^13^C]glycine for 2.5 h and tissue and serum glycine and acyl-glycine labeling were measured. (B) Circulating acyl-glycines are made from circulating glycine. Mean ± s.e. N = 4 mice. (C) Tissue phenylpropionylglycine labeling (normalized to circulating glycine labeling). Mean ± s.e. N = 4 mice. (D) Tissue butyrylglycine labeling (normalized to circulating glycine labeling). Mean ± s.e. N = 4 mice.

**Figure S5.**
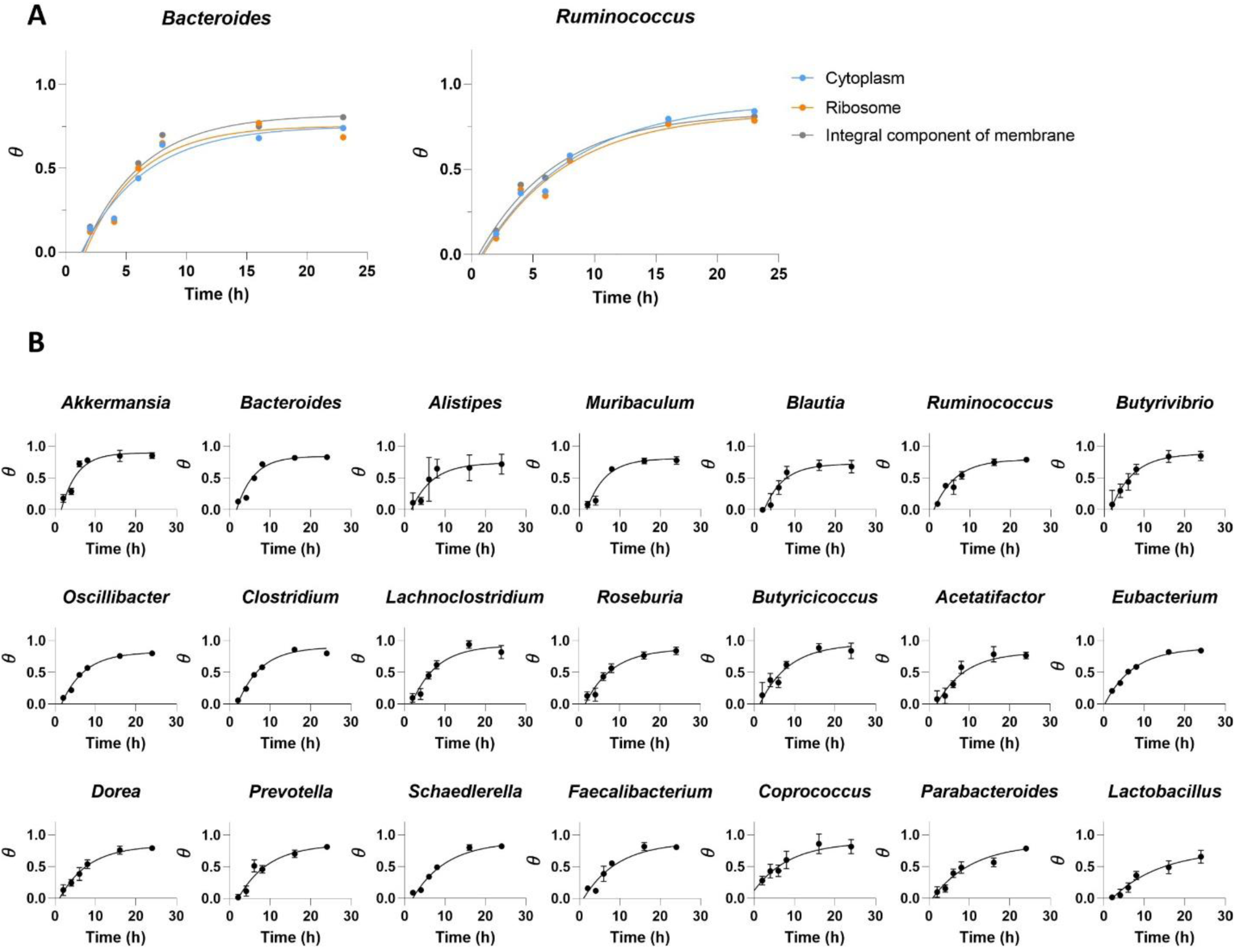
Single exponential fit of newly synthesized fraction of microbial peptides over time. (A) Different cellular compartments from the same bacterial genus show similar labeling rate. (B) Single exponential fit was applied to determine genus-level microbial turnover. Data are mean ± s.e. N = 5 mice.

**Figure S6.**
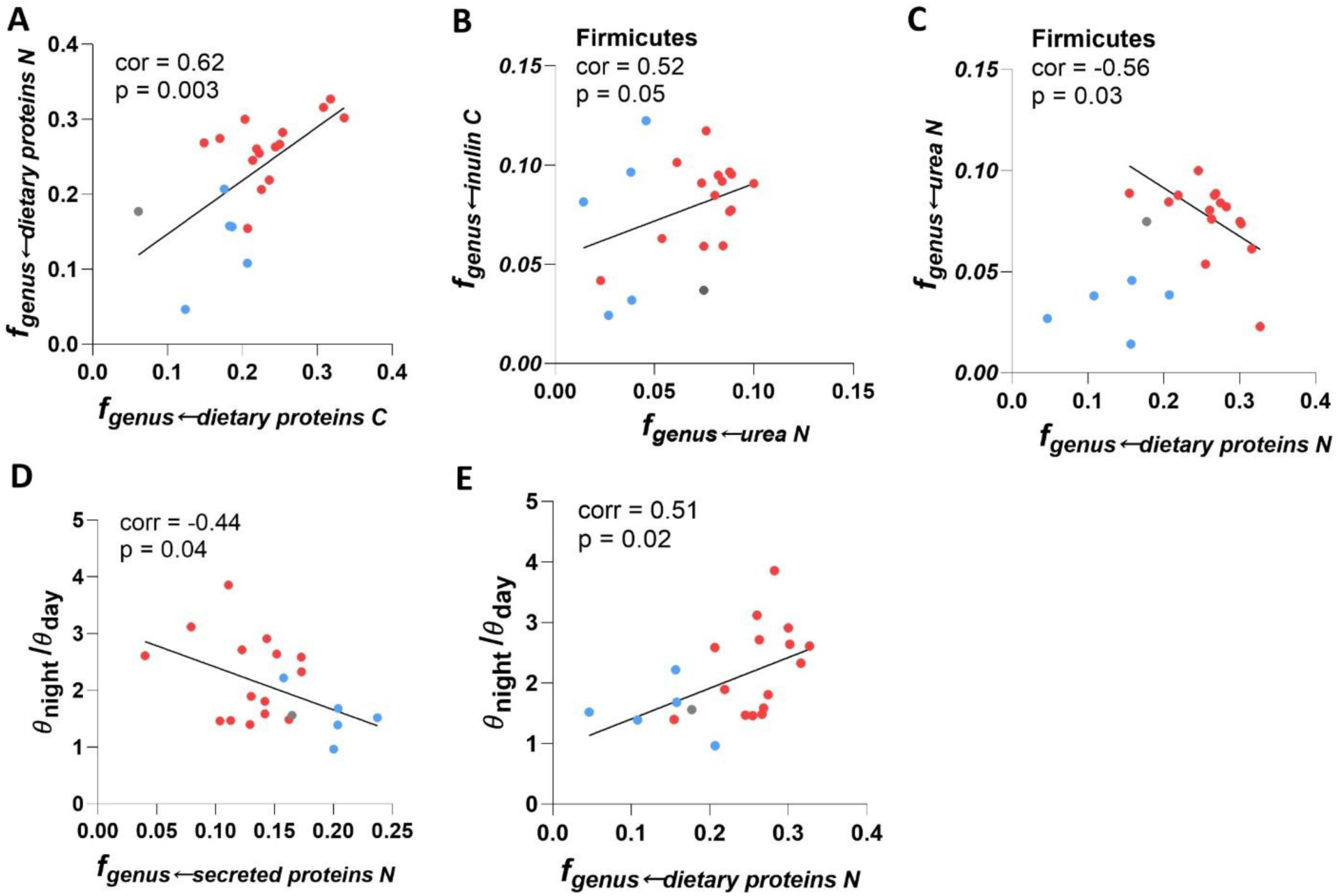
Correlation analysis of nutrient preferences across different gut bacterial genera. (A) Positive correlation between *f*_*genus*←*dietary proteins C*_ and *f*_*genus*←*dietary proteins N*_. (B) Positive correlation between *f*_*genus*←*inulin C*_ and *f*_*genus*←*urea N*_ in Firmicutes. (C) Negative correlation between *f*_*genus*←*urea N*_ and *f*_*genus*←*dietary proteins N*_ in Firmicutes. Negative correlation between *θ*_*night*_/ *θ*_*day*_ and *f*_*genus*←*secreted proteins N*_. Positive correlation between *θ*_*night*_/ *θ*_*day*_ and *f*_*genus*←*dietary proteins N*_.

**Figure S7.**
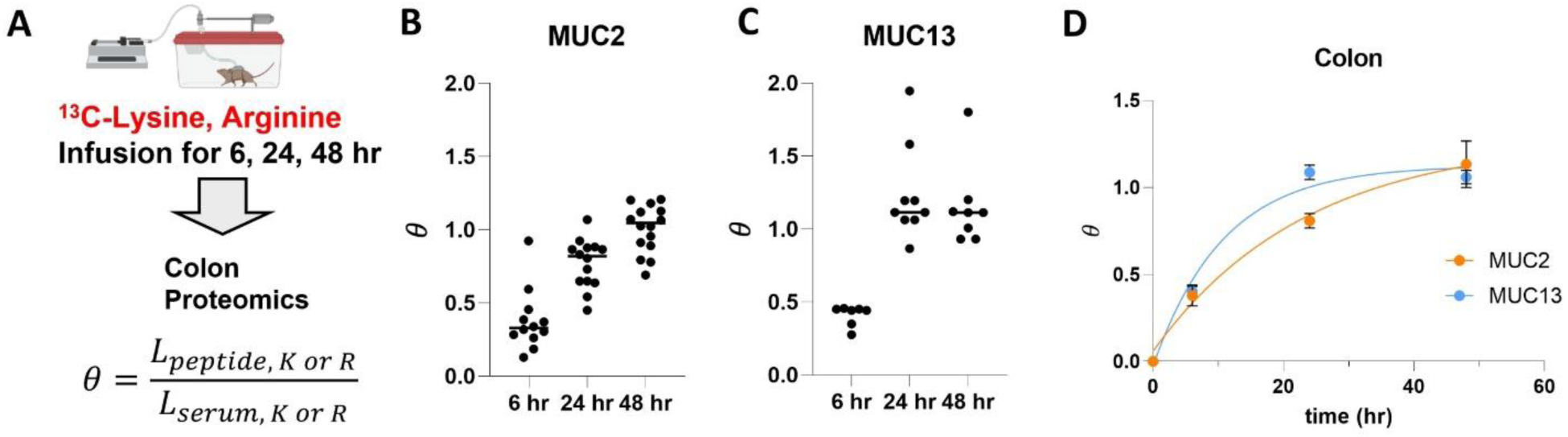
Host secreted proteins are synthesized from host circulating amino acids. (A) Experimental design. Mice were infused with ^13^C-lysine and ^13^C-arginine and mouse colon protein was analyzed by proteomics. (B) MUC2 labeling (multiple different MUC2 peptides). N = 3 mice. (C) MUC13 labeling (multiple different MUC13 peptides). N = 3 mice. (D) Single exponential fit of labeling fraction of MUC2 and MUC13 over time. Mean±s.e. N = 3 mice.

